# A novel workflow for unbiased quantification of autophagosomes in 3D in *Arabidopsis thaliana* roots

**DOI:** 10.1101/2023.09.11.557125

**Authors:** Michal Daněk, Daniela Kocourková, Tereza Podmanická, Kateřina Eliášová, Kristýna Nesvadbová, Pavel Krupař, Jan Martinec

**Affiliations:** Czech Academy of Sciences, Institute of Experimental Botany, Rozvojová 263, 165 00 Prague, Czech Republic

**Keywords:** ATG8, Cavalieri principle, differentiation zone, image analysis, immunofluorescence, meristematic zone, microscopy, optical disector, stereology

## Abstract

Macroautophagy is frequently quantified by live imaging of autophagosomes decorated with a marker of fluorescently tagged ATG8 protein (FT-ATG8) in *Arabidopsis thaliana*. This requires generation of suitable plant material by time-consuming crossing or transformation with FT-ATG8 marker. Autophagosome quantification by image analysis often relies on their counting in individual focal planes. This approach is prone to deliver biased results due to inappropriate sampling of the regions of interest in the Z-direction, as the actual 3D distribution of autophagosomes is usually not taken into account. To overcome such drawbacks, we have developed and tested a workflow consisting of immunofluorescence microscopy of autophagosomes labelled with anti-ATG8 antibody followed by stereological image analysis employing the optical disector and the Cavalieri principle. Our immunolabelling protocol specifically recognized autophagosomes in epidermal cells of *A. thaliana* root. Higher numbers of immunolabelled autophagosomes were observed when compared with those recognized with FT-*At*ATG8e marker, suggesting that single *At*ATG8 isoform markers cannot detect all autophagosomes in a cell. Therefore, immunolabelling provides more precise information as the anti-ATG8 antibody recognizes virtually all *At*ATG8 isoforms. The number of autophagosomes per tissue volume determined by stereological methods correlated with the intensity of autophagy induction treatment. Compared to autophagosome quantifications in maximum intensity projections, stereological methods detected autophagosomes present in a given volume with higher accuracy. Our novel application of immunolabelling combined with stereological methods constitutes a powerful toolbox for unbiased and reproducible quantification of autophagosomes and offers a convenient alternative to the standard of live imaging using FP-ATG8 marker.

## INTRODUCTION

Autophagy is a complex of essential biological processes providing reutilization and recycling of molecules and structures within a cell. Macroautophagy is one of these processes, characterized by the presence of a specific double membrane compartment called the autophagosome, in which the material destined for degradation in the vacuole is physically isolated from the surrounding cytoplasm [1]. In this article, macroautophagy will hereinafter be referred to as autophagy. Autophagy is controlled by a precisely coordinated network of expression of autophagy-related genes (ATG). ATG8 protein is widely used as a marker of autophagosomes as it is present at the autophagosome membrane from the stage of the phagophore [2] up to the degradation in the vacuole. Quantification of autophagosomes in a cell or tissue comprises the main method for studying autophagy intensity in plants [3,4]. Although there are reports on staining autophagosomes in plant cells with fluorescent dyes commonly used to detect autophagosomes in Metazoan cells [3,5], their specificity for autophagosomes in *Arabidopsis thaliana* roots has been questioned [6]. Genetically encoded ATG8 tagged with a fluorescent protein (FP-ATG8) thus remains a gold standard marker used for the identification and quantification of autophagosomes in plant science.

In *A. thaliana*, a model organism in plant biology, nine isoforms of ATG8 (*At*ATG8a – *At*ATG8i) were identified [7]. *At*ATG8a [8–11], *At*ATG8e [3,6,8,12–14] or *At*ATG8f [2,8,15] are commonly used as autophagosome FT-ATG8 markers in live imaging. The use of FP-ATG8 also carries some peculiarities, as overexpression of individual *At*ATG8 isoforms enhances the autophagy rate in general and increases the number of detected autophagosomes [16]. Moreover, different isoforms used in FP-ATG8 detected variable amounts of autophagosomes [8]. The variability of constructs used for FP-ATG8 expression in the scientific community may reduce the reproducibility of results among different research teams or even decrease the plausibility of research outcomes.

Apart from the mentioned challenges, the FP-ATG8 marker approach requires the introduction of the marker to the plant material of interest by crossing or transformation. The time-consuming generation of suitable lines can be inconvenient, e. g. in the case of screening for the autophagy-deficient phenotype in a large collection of mutants. To circumvent these hurdles, we decided to visualize autophagosomes by immunofluorescence imaging using an anti-ATG8 antibody. To our best knowledge, immunofluorescence imaging of autophagosomes has only been used a few times in plants [17–19].

Quantification of cellular structures often relies on the analyses of individual images. When confocal microscopy is employed, these images correspond to thin optical sections. Such an approach is also widely used to count autophagosomes [2,8]. However, autophagosomes are localized throughout the whole 3D volume of the cytoplasm. Quantification in individual planes represents an unacceptable simplification of the actual autophagosome localization and an underestimation of the variability of their abundance in the Z-direction. This may lead to incomplete or even misleading results on autophagosome counts. A toolbox of stereological methods has been developed for appropriate sampling of a cell or a tissue in 3D to provide an unbiased estimate of many quantifiable features, such as the number, volume, length or surface of cellular objects [20,21]. Discrete particles distributed within a cell or tissue, such as autophagosomes, can be easily counted using the disector method. A disector is a 3D sampling body (usually in the shape of a cube or orthogonal cuboid) inserted into the tissue under investigation. The objects are counted inside this body [22]. A variant of this approach, called optical disector, uses optical sectioning of experimental material, which can be achieved, for example, by taking Z-stacks with a confocal microscope [23]. Intriguingly, the need for unbiased quantification of autophagosomes by stereological methods has been articulated [24]. We intended to establish a novel workflow for stereological quantification of autophagosomes visualized by immunofluorescence using anti-ATG8 antibody in *A. thaliana* roots.

Our immunolabelling protocol specifically recognized autophagosomes in the epidermal cells of the root differentiation (DZ) and meristematic zone (MZ). Using the optical disector for autophagosome counting and the Cavalieri principle to estimate tissue volume we determined significant increase in number of autophagosomes reflecting the intensity of autophagy. More autophagosomes were identified by immunolabelling than with the FP-*At*ATG8e marker, suggesting that *At*ATG8e does not detect all autophagosomes in a cell. Immunolabelling thus provides more accurate information about the total number of autophagosomes, as the anti-ATG8 antibody recognizes virtually all *At*ATG8 isoforms. Combination of immunolabelling of autophagosomes and their subsequent stereological quantification offers a powerful toolbox for unbiased and reproducible quantification of autophagosomes.

## METHODS

### Plant material and autophagy induction

*Arabidopsis thaliana* Col-0 ecotype was used as wild type, and all mutant and marker lines were of Col-0 background. T-DNA insertional line *atg10-1* SALK_084434 and autophagosome marker lines expressing 35S::GFP-ATG8e and 35S::mCherry-ATG8e are described elsewhere [13,25]. 35S::GFP-*At*ATG8a line was purchased from NASC (stock code: N39996). Double labelled line 35S::mCherry-ATG8e × 35S::GFP-*At*ATG8a was generated by crossing the parental lines, and F1 generation of the progeny was used in experiments. Seeds were surface sterilized with commercial bleach and stratified at 4 °C for two days. They were sown onto 1% agar (Sigma) plates containing half-strength Murashige and Skoog salts (Caisson) supplemented with 1% sucrose and grown for six days in vertical position at 16 hours of light (22 °C, light intensity: 100 µmol · m ^-2^ · s ^-1^ PAR) 8 hours of darkness (20 °C). For the control treatment, seedlings were transplanted onto plates with the medium of the same composition as before, and the plates were placed in the same light conditions. For the autophagy induction treatment, seedlings were transplanted onto 1% agar plates containing half-strength Murashige and Skoog salts without nitrogen (Caisson), sucrose was omitted from the medium, and the plates were placed in the dark. The duration of the treatment was 2 or 6 hours.

### Immunolabelling

The immunolabelling protocol was based on a procedure used for *A. thaliana* wholemount samples [26]; some modifications based on [27] were introduced. Briefly, microtubule-stabilizing buffer (MTSB) was used throughout the procedure. Roots of treated seedlings were cut and put in incubation baskets (Intavis) for fixation with 4 % formaldehyde in MTSB + 0.1 % TritonX-100 (MP Biomedicals) for 1 h. The subsequent immunolabelling procedure, including cell wall digestion, membrane permeabilization, incubation with antibodies and final counter-staining with CalcoFluor White M2R (Sigma), was run using automated pipetting station *Insitu*Pro VS (Intavis, Biological Instruments AG). Rabbit anti-ATG8 (Agrisera) and anti-rabbit conjugated to DyLight488 (Agrisera) or AlexaFluor555 (Invitrogen) were used as primary and secondary antibodies, respectively. For detailed protocol and recipes, consult Supplement 1.

### Microscopy, stereological quantification and co-localization assessment

Immunolabelled roots were mounted in 50% glycerol containing 0.02 % sodium azide. Strips of one layer of parafilm were used as spacers between a microscopic slide and a coverslip to prevent mechanical damage to roots. For live imaging, whole seedlings were mounted in a drop of liquid medium of the same composition as the medium used for a given treatment. A spinning disk microscope (Nikon Eclipse Ti-E, with Yokogawa CSU-W1 spinning disk unit) equipped with Prime BSI sCMOS camera (Photometrics) was used for Z-stack acquisition. Z-step was set to 300 or 500 nm. Water immersion 40× objective (Nikon, N.A. 1.25) was used. The total number of collected steps within a Z-stack was always kept higher than the minimal number (i.e., 21 for Z-step = 500 nm or 35 for Z-step = 300 nm) to cover 7.5 µm in the Z-direction, which was the Z-dimension of the disector we used for stereological quantification. Fluorescence signals were collected using the following settings. GFP or DyLight488: 488 nm laser excitation and 525/30 nm filter cube. mCherry of Alexa555: 561 nm laser excitation and 641/75 nm filter cube. CalcoFluor White: 405 nm laser excitation and 447/60 nm filter cube. Omicron LightHUB Ultra Lasers and Semrock Brightline filter cubes were used. Sequential acquisition was performed in the case of multi-channel imaging.

Autophagosome counting was performed by applying the optical disector method [22,23]. Individual autophagosomes were counted by clicking using the point tool in Fiji [28] with an installed Disector plugin (https://imagej.nih.gov/ij/plugins/sampling-window/index.html). Single cells were not distinguished; all the cells within the disector were analysed together. Different settings for disector in DZ and MZ were applied. In MZ, two disectors of (X; Y; Z) dimensions equal to (50; 50; 7.5) µm were placed in the epidermal layer. The volume of one disector in MZ was thus equal to 18750 µm^3^. The estimated number of autophagosomes per volume of tissue *estNv* in MZ was then calculated as follows:

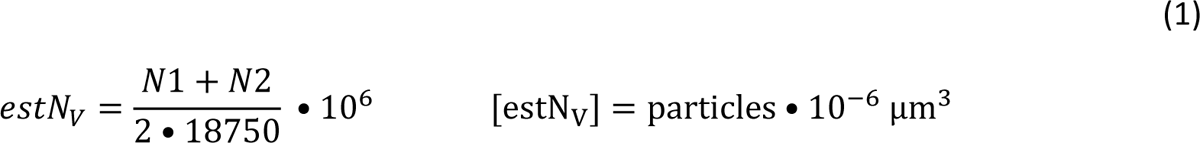

where *N1* and *N2* are the numbers of autophagosomes counted within the first and second disector

In DZ, one larger disector of (x; y; z) dimensions equal to (150; 150; 7,5) µm was used (Fig. 1). Such an approach was chosen because of the relatively lower density of autophagosomes in this tissue and quite a high variation in the number of particles among individual cells. The volume of disector intersecting the root was estimated using the Cavalieri principle and point counting [20,29] using the Grid tool in Fiji [28] according to:

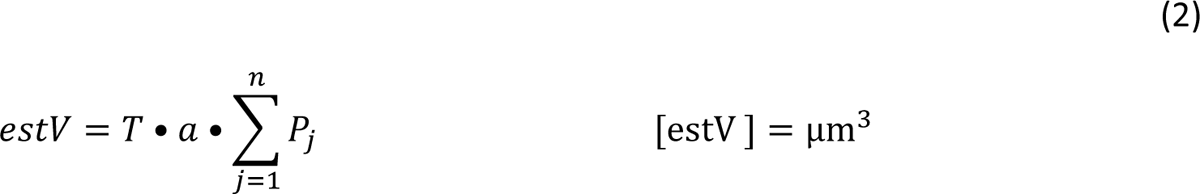

where *estV* is the estimator of volume in µm^3^, *T* is the distance between two neighbouring optical sections, *a* is the area in µm^2^ represented by one testing point of the testing grid, and *Pj* is the number of testing points hitting the j-th (j = 1, 2, …, *n*) optical section. In our case, *a* was 300 µm^2^, and T was 1.5 µm. The estimated number of autophagosomes per volume of tissue *estNv* [30] in DZ was then calculated as follows:

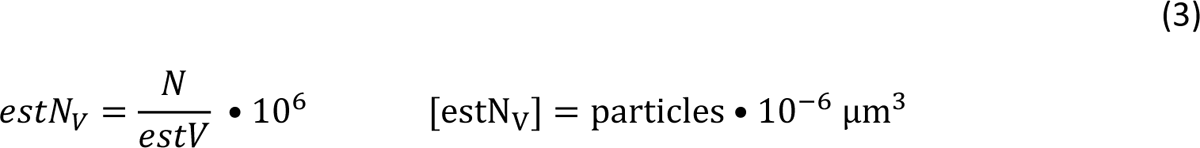

where *N* is the number of autophagosomes counted within the disector, and *estV* is the estimator of volume in µm^3^ determined in Equation (2).

Attention was paid to put disectors correctly in the given root zones. As there were particles of high fluorescence adhering to the root surface, probably produced through the immunolabelling procedure, and these particles could be mixed up with autophagosomes, we first checked that the position of the first plane of the disector was below the root surface. The signal of CalcoFluor White staining cell walls was observed for this purpose and to verify that the cells did not change shape or were not damaged during the fixation and immunolabelling process.

Co-localization analyses were carried out by counting particles in two channels separately and by verifying the presence of the signal in either of them using optical disector as described above.

**Figure 1:**
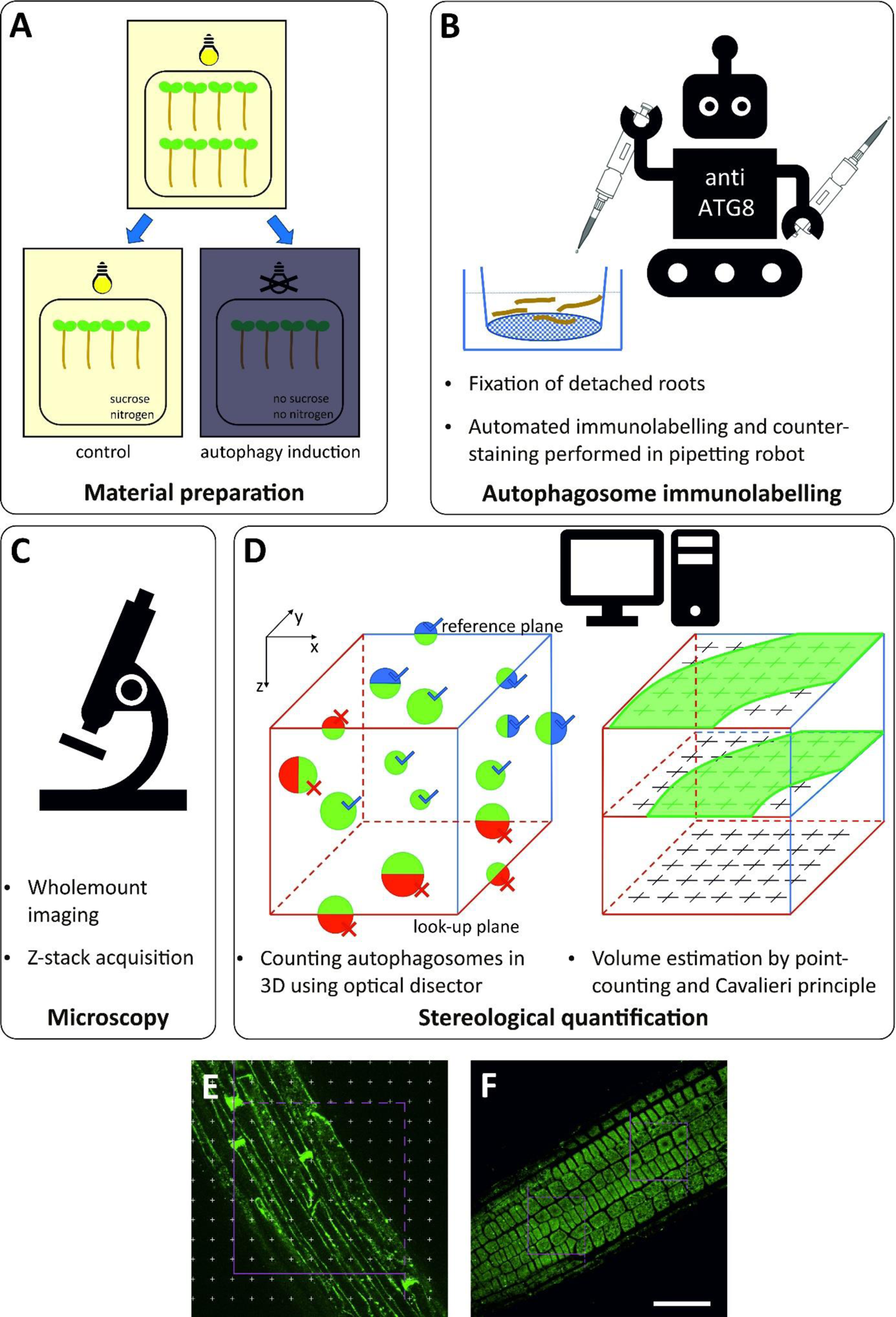
Workflow of stereological quantification of immunolabelled autophagosomes. **A**: Seedlings are grown for six days in the control medium, autophagy is induced by carbon and nitrogen starvation for several hours. **B**: Detached roots are placed in incubation baskets where they are fixed and subsequently undergo immunolabelling protocol and staining with CalcoFluor White performed by a pipetting robot. **C**: Samples are mounted in 50% glycerol and observed using confocal microscopy. Z-Stacks are acquired for stereological quantification. **D:** Quantification is a two-step process. First, individual particles are counted using optical disector. Optical disector is a virtual prism delimiting the volume where individual particles are counted. In the Z-direction, it is restricted by two focal planes. Particles apparent at the reference plane as well as all the particles that appear in the volume of disector are included in the final number of autophagosomes under the condition that they are not apparent at the look-up plane (green-blue balls at the top and green ball inside vs green-red balls at the bottom of the disector). In X, Y dimension two types of edges delimit the disector. Particles present inside the disector that intersect exclusion edges (red lines) are excluded (green-red balls at sides) whereas particles inside the disector that intersect inclusion edges (blue lines) are included (green-blue balls at sides of the disector). Particles within the disector that do not intersect any line or plane are included (green balls). If the tissue of interest does not occupy the whole volume of the disector it is necessary to estimate the volume of tissue inside the disector. For this purpose, Cavalieri principle combined with point-counting method is applied. A grid with evenly distributed points each of which represents defined surface area is superimposed on several Z-planes spaced by the same known distance in the Z-direction within the disector. Projections of tissue of interest at these Z-planes are shown as green areas. Each point of the grid thus represents a known unit of volume. The points hitting the tissue of interest are counted (green crosses), and the volume of the tissue intersecting the disector is estimated. **E, F**: Immunolabelled root epidermal cells of the differentiation (**E**) and meristematic (**F**) zone with superimposed disector (magenta lines, the reference plane is shown) used in our analysis. **E**: Disector (x; y) dimensions equal to (150; 150) µm were used. At these dimensions, the root does not fill the whole disector and subsequent estimation of the volume of the root intersected by the disector is required. The point grid (white crosses) is used for that purpose. **F**: Two disectors of (x; y) dimensions equal to (50; 50) µm were placed in the meristematic zone. Both disectors are fully filled with the root so additional estimation of the sampled root tissue volume is not necessary. Scale bar: 50 µm.

Maximum intensity projections were constructed in Fiji to present data more illustratively. Image processing consisted in contrast and brightness adjustment; images presented as parts of the Figures were rotated when necessary. All the processing and analyses were conducted in Fiji.

### Statistical evaluation

Statistical tests were carried out in MS Excel (Microsoft) and Prism 8 (GraphPad Software). In each data set, outliers were excluded from statistical analysis. Outliers were considered values exceeding 1.5-multiple of interquartile range. Plots were built up in Prism 8. Figures were assembled in Corel DrawX7 (Corel Corporation).

### Transient expression of AtATG8 isoforms in Nicotiana benthamiana leaves

*Agrobacterium tumefaciens* strains harbouring pUBQ10::*At*ATG8a-i [9] (courtesy of Dr Yasin Dagdas, Gregor Mendel Institute of Molecular Plant Biology, Vienna, Austria) or p19 viral suppressor of silencing [31] were grown in liquid YEP medium (yeast extract: 10 g per litre, peptone: 10 g per litre, NaCl: 5 g per litre, pH = 7.0) overnight. Cultures were rinsed and resuspended in infiltration buffer (10 mM MgSO4, 10 mM MES, pH = 5.6). Final OD600 was 0.1 for GFP-*At*ATG8a-i, and 0.05 for p19. Leaves of four-week-old *N. benthamiana* plants grown in 16 hours light (22 °C, light intensity: 100 µmol m ^-2^ ·s ^-1^ PAR), 8 hours dark (20 °C) regimen were agroinfiltrated with a syringe and kept in low light conditions for 16 hours. Then they were moved to their previous conditions. Fluorescence was observed approx. 40 hours after infiltration.

### Total protein extraction and Western blot analysis

Approx. 80 mg of agroinfiltrated *N. benthamiana* leaf tissue was frozen in liquid nitrogen and homogenized with beads in an Eppendorf tube for 3 mins at f = 20 Hz using a tissue grinder (Retsch). Ice-cold extraction buffer (50 mM Tris-HCl, pH = 7.5, 150 mM NaCl, 10% glycerol, 2 mM EDTA, 50 µM DTT, 0.5% Igepal) with cOmplete Protease Inhibitor Cocktail (Roche) was added to homogenized samples (500 µL buffer per 100 mg fresh weight), the mixture was vortexed and kept on ice for 30 minutes. Samples were centrifugated at 5 000 g for 5 min at 4°C to remove debris. Supernatants were collected, and total protein concentration was determined by Coomassie Plus Protein Assay Reagent (Thermo Scientific) in a plate reader. Samples with a total protein concentration of 2 µg per µL were prepared and denatured at 95 °C for 10 mins. Proteins were separated on 10% SDS-PAGE and blotted onto nitrocellulose membranes (Serva) by wet transfer. Membranes were blocked in 5% low-fat milk in TBS-T for 2 hours and probed with 1:5000-diluted mouse anti-GFP (Roche) or rabbit anti-ATG8 (Agrisera) antibody in 3% low-fat milk in TBS-T for 2 hours. Three consecutive washes with TBS-T, each for 15 mins, were carried out prior to the incubation with secondary 1:10 000-diluted anti-rabbit (Bethyl Laboratories) or anti-mouse (Promega) in 5% low-fat milk in TBS-T for 1 hour. Three consecutive washes with TBS-T, each for 15 mins, followed by one wash with TBS for 5 mins, were done. Membranes were developed with chemiluminescent substrate (Azure Biosystems) and imaged using Sapphire Biomolecular Imager (Azure Biosystems). Band intensities were measured using the Gels tool in Fiji. Attention was paid to analyse undersaturated images.

## RESULTS

To provide a robust induction of autophagy in roots, we chose a widely applied treatment of nitrogen and carbon starvation [4,8] in 6-day-old seedlings on a solid medium (Fig. 1 A). Autophagosomes are usually quantified in the root DZ. This preference is based on the low number of autophagosomes in this zone under control conditions, especially when compared to MZ. An increase in autophagosome number is thus apparent upon autophagy-inducing treatment [4]. On the other hand, the immunofluorescence approach in Arabidopsis root is applied rather to visualize structures in MZ; its use in DZ is relatively scarce (e.g. [32,33]). We decided to investigate and compare the possibility of autophagosome immunolabelling in both zones throughout our study.

First, we employed an established immunolabelling protocol for *A. thaliana* roots routinely used to observe cytoskeleton or membrane proteins [26]. However, we had to adjust the procedures since many DZ epidermal cells collapsed when the original protocol was used. Such damage might be caused by the high osmolarity of the solutions used in the protocol, mainly by the presence of mannitol in the cell wall digestion step. Hence, we omitted mannitol from the solution and used 0.1% Triton X-100 already in the fixation step, which enabled better penetration of the fixative through the root tissue [27]. We also introduced a step of counter-staining cell walls with CalcoFluor White following the immunolabelling procedure. The introduction of this step helped us to assess the state and shape of cells and correctly position the disector in the tissue. The adjusted protocol produced better results, with most labelled roots being undamaged in DZ (Fig. 1 B). The protocol is available in detail in Supplement 1. Z-stacks were obtained using a spinning disk microscope (Fig. 1 C), allowing fast acquisitions of several tens of Z-planes within a Z-stack. Disector for stereological quantification (Fig. 1 D) was set differently in root DZ and MZ (Fig. 1 E, F).

To investigate the feasibility of immunolabelling for autophagosome detection, we induced autophagy for 2 hours. In Col-0 seedlings, we observed a few particles resembling autophagosomes already in control conditions in both root zones investigated (Fig. 2 A, D, G, K); however, the pattern was different in these zones. In DZ, the immunofluorescence signal associated with discrete particles was accompanied by a relatively high signal at the edges of cells; it was prominently strong at the apical and basal poles of the cells (Fig. 2 A). This signal probably represents the labelling of soluble non-lipidated ATG8 in the cytoplasm. In MZ, the diffuse signal from the cytoplasm was stronger than in DZ. Still, the signal from discrete particles was of much higher intensity making these particles easily detectable and separable from their surroundings (Fig. 2 D). The number of discrete particles increased upon autophagy induction in both zones (Fig. 2 B, E, H, L). In DZ, the diffuse signal in the cytoplasm decreased, whereas the signal associated with the particles increased (Fig. 1 B), probably at the expense of the former [2]. This could suggest the re-localization of the protein recognized by the anti-ATG8 antibody from the cytoplasm to the discrete particles that was incited by autophagy induction. To test the possibility of unspecific binding of the secondary antibody, we omitted the primary anti-ATG8 antibody from the immunolabelling protocol. In both DZ and MZ, only a very low signal was detected (Fig. 2 C, F). We included a mutant *atg10-1* reported to form virtually no autophagosomes [25] in our study to verify that the particles we detected using the immunolabelling procedure are associated with the autophagy process. No particles were observed in either root zone under control conditions or upon autophagy induction, while a relatively high diffuse signal was present in the cytoplasm (Fig. 2 I, J, M, N).

**Figure 2:**
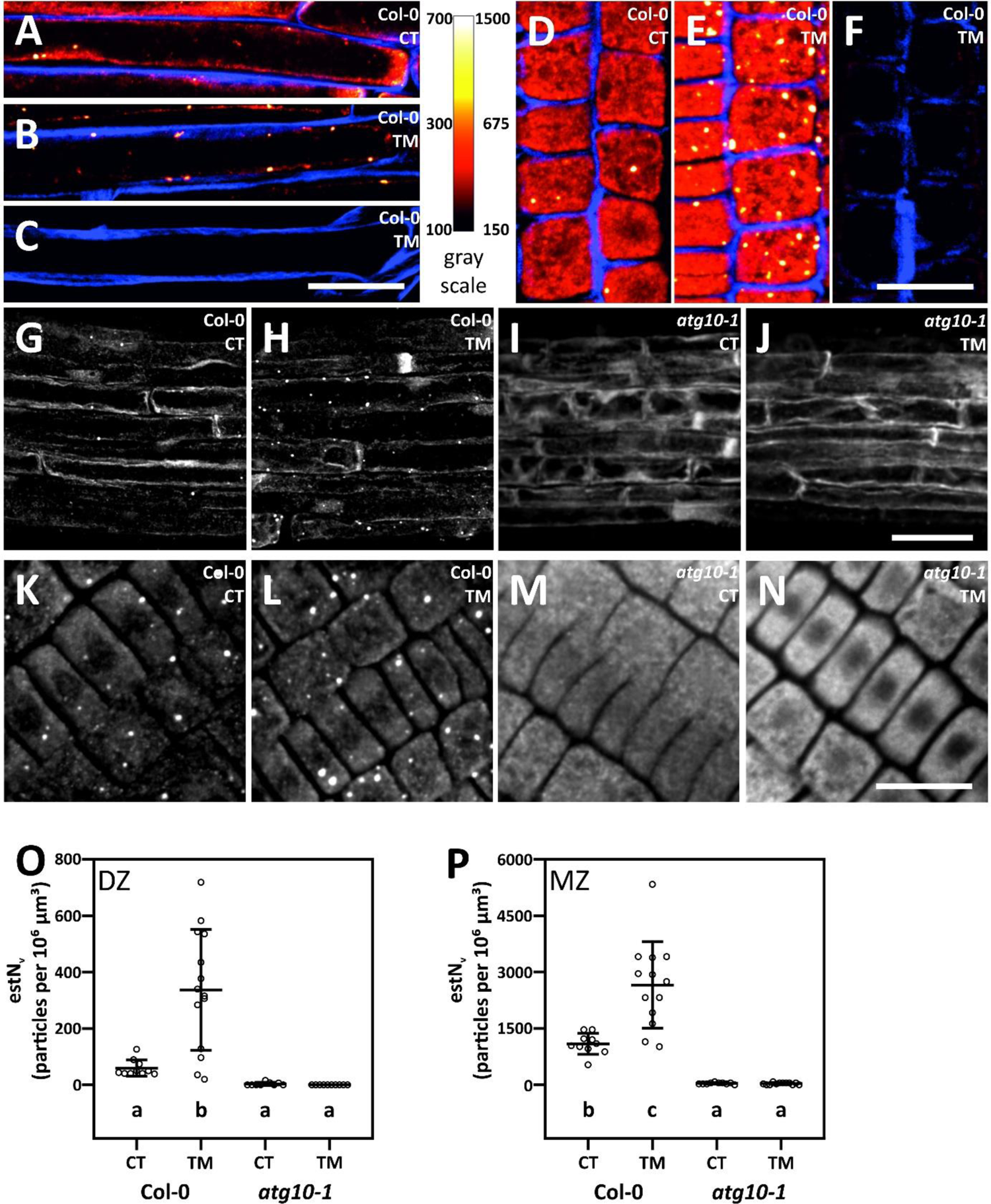
Anti-ATG8 antibody recognizes intracellular particles, density of which increases upon autophagy induction in Col-0. These particles are not formed in an autophagy-deficient mutant *atg10-1*. **A – F:** Detailed view at root epidermal cells in the differentiation (**A – C**) and meristematic (**D – F**) zone of Col-0 immunolabelled roots. **A, D**: seedlings grown in control conditions were transplanted to the same conditions for 2 hours (control, CT). **B, C, E, F:** seedlings grown in control conditions were transplanted to conditions inducing autophagy for 2 hours (treatment, TM). In **C, F** primary antibody was omitted from immunolabelling protocol. Immunofluorescence signal is presented in Red Hot LUT, cell walls were counter-stained with CalcoFluor White (in blue). Images represent maximum intensity projections of six Z-planes (i. e. 1.5 µm depth along the Z-axis). **G – N:** Maximum intensity projections of disector volumes used for stereological quantification in root epidermal cells in the differentiation (**G – J**) and meristematic (**K – N**) zone. **G, H, K, L:** Col-0. **I, J, M, N**: *atg10-1*. **G, I, K, M:** seedlings grown in control conditions were transplanted in control conditions for 2 hours. **H, L, J, N:** seedlings grown in control conditions were treated to induce autophagy for 2 hours. Roots of 6-day-old seedlings were immunolabelled, primary antibody: anti-ATG8, secondary antibody: anti-rabbit DyLight488. Spinning-disk microscope was used to acquire Z-stacks. Scale bars: 20 µm (**A – F, K – N**) and 50 µm (**G – J**). **O, P:** Quantification of number of particles per volume of root epidermal cells in the differentiation zone (DZ) and meristematic zone (MZ) in Col-0 and *atg10-1* in control and autophagy-inducing treatment. Means ± SD are presented. One-way ANOVA, letters indicate significant differences between groups by multiple comparison Tukey-Kramer test, p < 0.01, n = 6 – 14. Three biological replicates providing the same results were conducted independently.

Stereological quantification confirmed the increase in the number of particles per root epidermis volume in Col-0 in both zones after autophagy induction, as well as the absence of such a response in autophagosome formation-deficient mutant *atg10-1*, where no particles were detected regardless of conditions (Fig. 2 O, P). It is worth noting that the number of autophagosomes in MZ is roughly one order of magnitude higher than in DZ.

To verify the identity of detected particles as autophagosomes, we immunolabelled roots of lines expressing FP-*At*ATG8e autophagosome marker. We observed co-localization of immunolabelled particles and autophagosomes marked with FP-*At*ATG8e in both root zones. The number of co-localizing particles increased upon autophagy induction (Fig. 3 A – D, Fig. S1 A – D). Hence, we conclude that the immunolabelled particles are indeed autophagosomes and that our approach is suitable for autophagosome detection. Although the majority of autophagosomes showed dual labelling (i.e., co-localization), we also observed a minor population of particles with immunofluorescence signal that did not co-localize with autophagosomes labelled with GFP-*At*AT8e or mCherry-*A*tATG8e (Fig. 3 A – D, Fig. S1 A – D). When particles with immunofluorescence or fluorescent protein signal were quantified separately, higher numbers were gained for the former than for the latter (Fig. 3 E, F, Fig. S1 E, F). Accordingly, a significant percentage of autophagosomes was detected uniquely by immunofluorescence among all the autophagosomes observed while there was only a negligible population of autophagosomes labelled only with GFP-*At*ATG8e or mCherry-*At*ATG8e (Fig. S2 A, B). We suggest that anti-ATG8 antibody can recognize autophagosomes lacking *At*ATG8e, and such particles contribute to a significantly higher number of autophagosomes detected with immunofluorescence than with *At*ATG8 single-isoform based FP-ATG8 markers.

**Figure 3:**
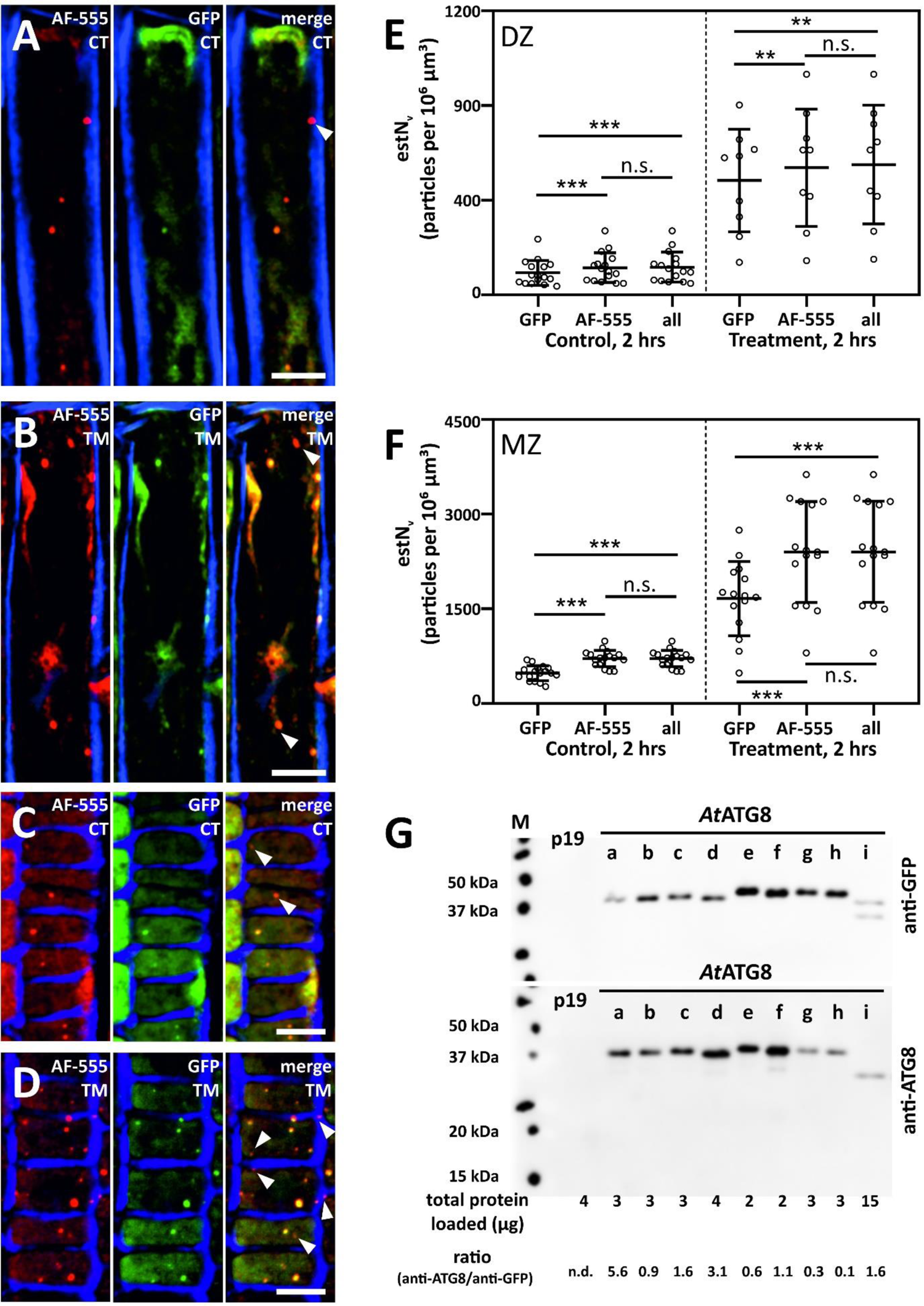
Anti-ATG8-labelled particles are autophagosomes. The immunolabelling approach allows the identification of a higher number of autophagosomes when compared to the quantification of autophagosomes labelled by a single isoform FP-ATG marker lines. **A – D:** Co-localization of immunolabelled particles (left panel, in red) with autophagosomes labelled with GFP-*At*ATG8e (middle panel, in green) in root epidermal cells in the differentiation zone (**A, B**) or meristematic zone (**C, D**) under control conditions (CT; **A, C**) or under treatment inducing autophagy (TM; **B, D)**. Note that some immunolabelled particles are not decorated with GFP-*At*ATG8e (arrowhead in merge images, right panels) whereas the majority of particles show clear signal in both channels. Roots of 6-day-old seedlings expressing 35S::GFP-ATG8e were immunolabelled, primary antibody: anti-ATG8, secondary antibody: anti-rabbit AlexaFluor555, and subsequently counter-stained with CalcoFluor White to visualize cell borders (in blue). Spinning-disk microscope was used to acquire Z-stacks. Maximum intensity projections of six Z-planes (i. e. 1.5 µm depth along the Z-axis) are presented. Scale bars: 10 µm. **E, F:** Stereological quantification of number of autophagosomes per volume of root epidermal cells in the differentiation (DZ) and meristematic zone (MZ). Particles were counted separately in individual channels (GFP: sum of particles detected with GFP-*At*ATG8e; AF-555: sum of particles detected with anti-ATG8/anti-rabbit AlexaFluor555; all: sum of all detected particles). Means ± SD are presented. Pair-wise t-test, ** p < 0.01; *** p < 0.001, n. s. not significant, n = 9 – 17. Three biological replicates providing the same results were conducted independently. **G:** Anti-ATG8 antibody recognizes all *At*ATG8 isoforms but with different affinity. Western blots of total protein extracts prepared from *N. benthamiana* leaves transiently expressing individual isoforms of GFP-*At*ATG8. Transformation with p19 only was used as negative control. Two gels were loaded with the same amounts of total proteins for each extract. The upper and lower membrane were probed with anti-GFP antibody and anti-ATG8 primary antibody, respectively. Compare different ratios of signal intensities between bands labelled with anti-ATG8 and anti-GFP for different *At*ATG8 isoforms. Note that untagged *Nt*ATG8 isoforms were not detected (molecular mass ca. 15 kDa). Molecular mass in kDa for *At*ATG8a-i tagged with GFP follow: 42.4; 40.9; 40.7; 45.9; 40.9; 40.8; 40.9; 40.9; 40.3.

As the anti-ATG8 antibody was described by the manufacturer to recognize multiple *At*ATG8 isoforms, we wanted to verify how efficient the antibody is to all of them. We successfully expressed GFP-*At*ATG8a-i single isoforms in *Nicotiana benthamiana* leaves. Under control conditions, all nine isoforms were localized in the cytoplasm with a few autophagosomes (Fig. S3 A – I) resembling the situation observed in *A. thaliana* root epidermal cells in control conditions. Protein extracts isolated from transformed leaves were tested for anti-ATG8 reactivity by Western blot analysis. Clear bands corresponding with the expected mass of GFP-*At*ATG8 were observed for all the isoforms except *At*ATG8i, where a lighter band occurred when anti-ATG8 or anti-GFP primary antibodies were used. Bands in GFP-*At*ATG8i lanes were detected upon high sample loading, suggesting that the observed bands are not specific for *At*ATG8i (Fig. 3 G). To assess the affinity of anti-ATG8 antibody to the individual isoforms we compared the intensities of bands acquired by probing the membrane with anti-ATG8 antibody with the ones where fusion proteins were detected by anti-GFP antibody. Similar or higher ratios than for *At*ATG8e were reached for six other *At*ATG8 isoforms, out of which *At*ATG8a, *At*ATG8b, *At*ATG8d, *At*ATG8f and *At*ATG8i are expressed in roots of 6-day old seedlings [34]. These isoforms can decorate the immunolabelled autophagosomes devoid of the *At*ATG8e marker observed in our co-localization analysis.

To test whether sub-populations of autophagosomes exist that are differentially decorated by specific *At*ATG8 isoforms, we crossed *A. thaliana* marker lines expressing GFP-*At*ATG8a and mCherry-*At*ATG8e, which are among the most often used autophagosome markers, and assessed co-localization of autophagosomes labelled with both isoforms in DZ (Fig. S3 K, L). While most of the autophagosomes detected under control and autophagy-inducing conditions were labelled with both isoforms, autophagosomes decorated uniquely by one or the other isoforms occurred concomitantly (Fig. S3 K – N).

Taken together, our findings demonstrate that immunolabelling using anti-ATG8 recognized more autophagosomes than FP-*At*ATG8 single isoform markers, probably due to binding to autophagosomes decorated by virtually all *A*tATG8 isoforms expressed in a given tissue.

We have already demonstrated an increase in the number of autophagosomes from control conditions to autophagy induction. As a next step of our analysis, we tested whether our immunolabelling protocol coupled with stereological assessment of autophagosome number can unravel smaller differences in the number of autophagosomes, i. e. whether it is possible to detect gradual increase in the intensity of autophagy. In order to test that, we induced autophagy for two different periods of time. We supposed the 6-hour treatment to have a more pronounced impact on autophagosome formation than the 2-hour treatment we have used in our study so far. First, we tested this setup using live imaging of autophagosome marker line expressing GFP-*At*ATG8e that we used as a benchmark, to see whether the expected difference in autophagosome number depending on the duration of induction can be observed. As presumed, more autophagosomes were detected after 6-hour treatment than after 2 hours of autophagy induction in the GFP-*At*ATG8e marker line using live imaging (Fig. S4 C – E) as well as in immunolabelled Col-0 seedlings (Fig. 4 C, D, G – J). The same effect was found in both DZ and MZ in immunolabelled seedlings. We did not observe any difference between longer and shorter incubation in control conditions; (Fig. 4 A, B, E, F, I, J, Fig. S4 A, B, E). Immunolabelling followed by stereological quantification is thus able to distinguish more subtle changes in the autophagosome number and may be suitable, for example, to assess gradual levels of stressor severity or downregulation of endogenous pathways affecting autophagy.

**Figure 4:**
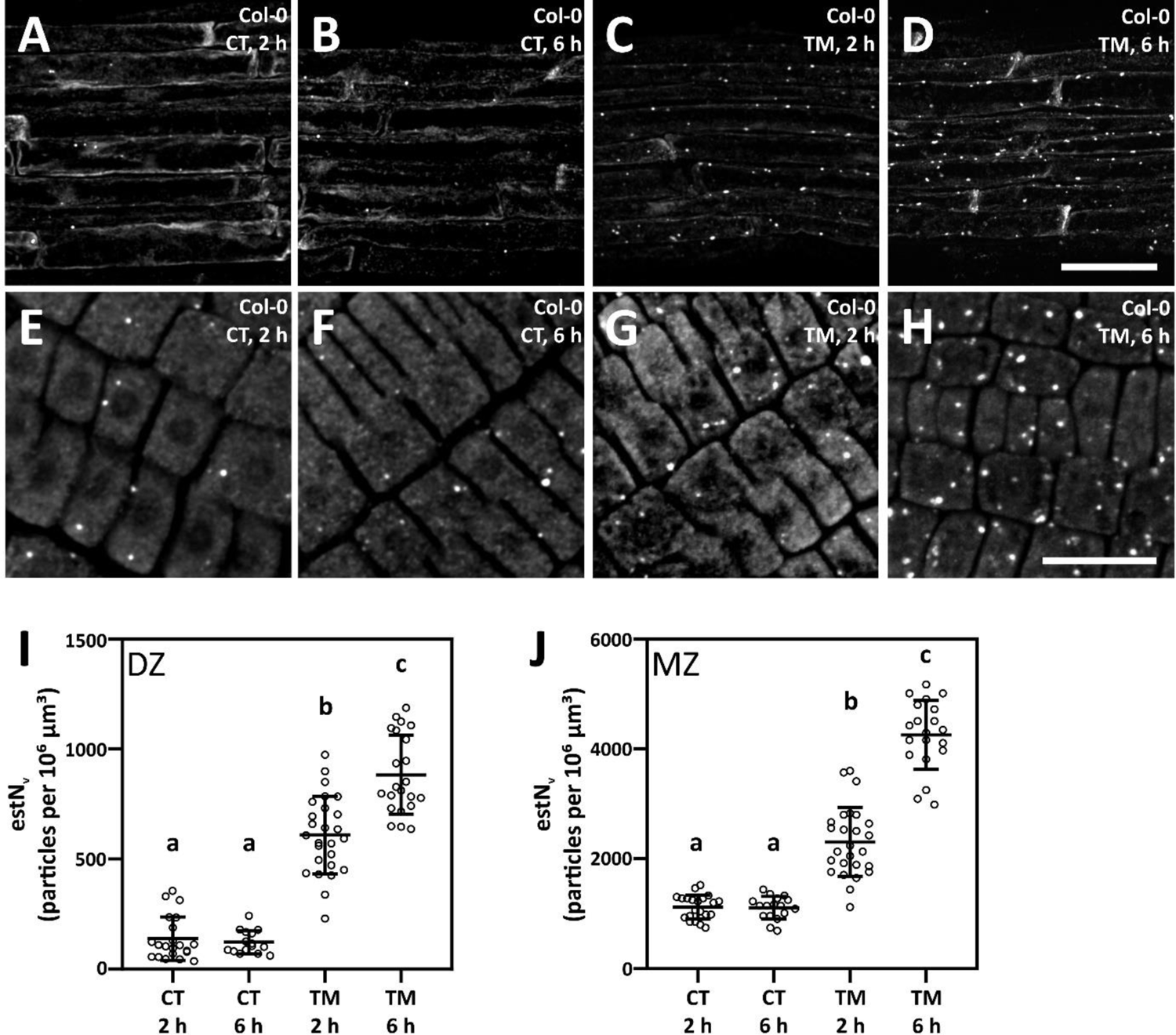
Immunolabelling with anti-ATG8 enables assessment of the intensity of autophagy. Stereological quantification of autophagosomes in root epidermal cells of differentiation (**A** – **D**) and meristematic zone (**E** – **H**) in Col-0. Seedlings grown in control conditions were transplanted to the same conditions (CT) for 2 hours (**A**, **E**) and 6 hours (**B**, **F**) or the conditions inducing autophagy (TM) for 2 hours (**C**, **G**) and 6 hours (**D**, **H**). Roots of 6-day-old seedlings were immunolabelled, primary antibody: anti-ATG8, secondary antibody: anti-rabbit DyLight488. Spinning-disk microscope was used to acquire Z-stacks. Images represent maximum intensity projections of disector volumes used for stereological quantification. Scale bars: 50 µm (**A – D**) and 20 µm (**E – H**). **I, J:** Quantification of number of particles per volume of root epidermal cells in the differentiation zone (DZ) and meristematic zone (MZ) in Col-0 in control and autophagy inducing treatment lasting for 2 hours and 6 hours. Means ± SD are presented. One-way ANOVA, letters indicate significant differences between groups by multiple comparison Tukey-Kramer test, p < 0.01, n = 15 – 26. Three biological replicates providing the same results were conducted independently.

Finally, we were interested in whether the stereological approach is not pointlessly complicated and whether we could gain the same results by analysing maximum intensity projection as in [17,35–37]. First, we counted autophagosomes using the optical disector as in the previous cases. We then used disectors as a region of interest where we constructed maximum intensity projection (MIP). The last focal planes of the disectors (i.e., look-up planes) were omitted from MIP construction. Finally, the autophagosomes were counted at MIPs. We observed consistent differences between quantification by disector and MIP; interestingly, the nature of the differences varied between DZ and MZ.

In DZ, the different results were present only when autophagy was induced, and more particles were detected at MIPs (Fig. 5 A). On the other hand, quantification using MIP gained fewer particles when compared with the disector in MZ but only under control conditions (Fig. 5 B).

**Figure 5:**
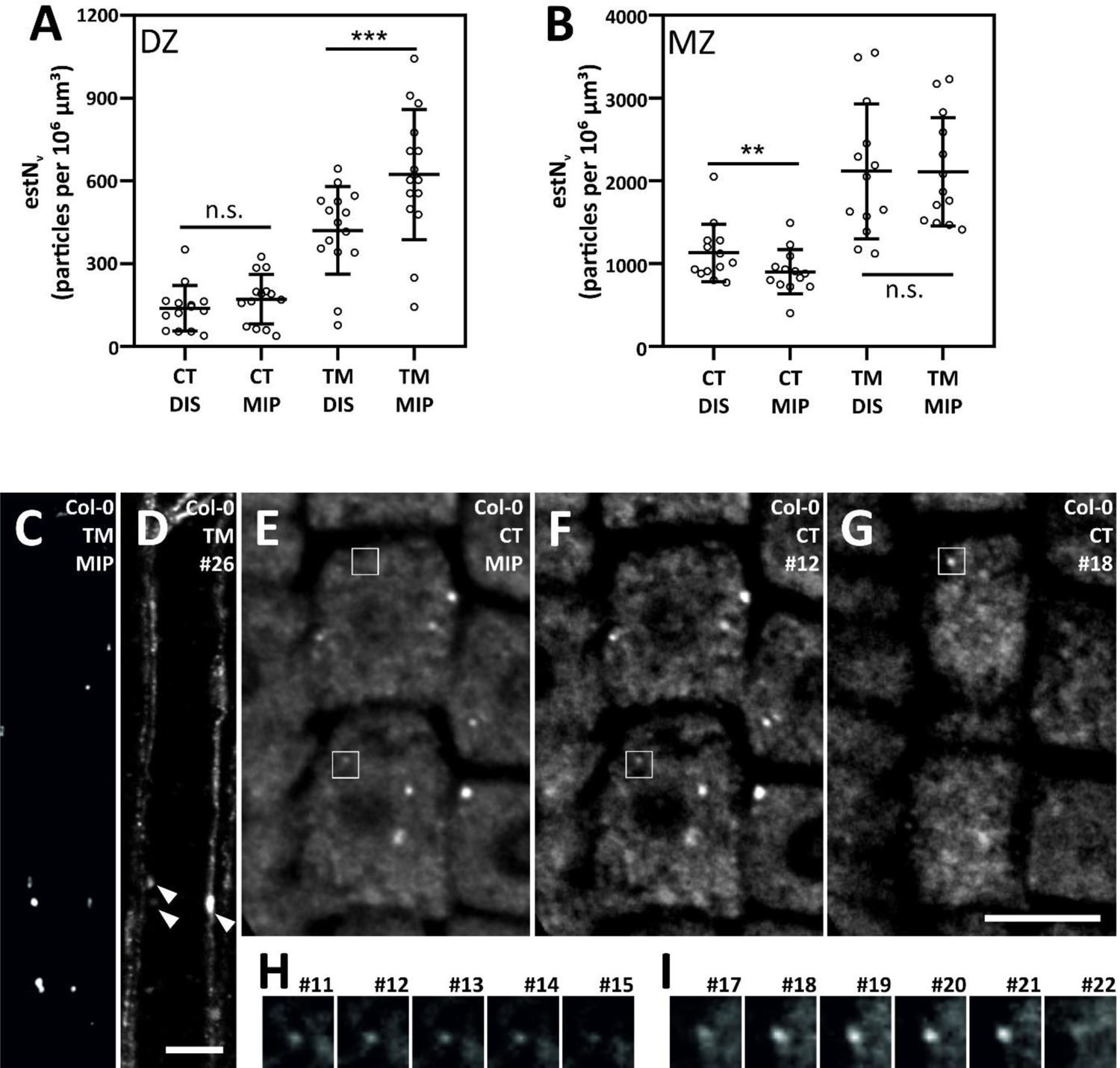
Stereological approach offers a more accurate assessment of autophagosome number than quantification using maximum intensity projection. **A, B**: Different results are obtained when autophagosomes are counted by using disector and at maximum intensity projections in differentiation zone (**A**) and meristematic zone (**B**) in Col-0 root epidermal cells. Six-day old seedlings were transplanted into control condition (CT) or treatment inducing autophagy (TM) for 2 hours and subsequently immunolabelled, primary antibody: anti-ATG8, secondary antibody: anti-rabbit DyLight488. Spinning-disk microscope was used to acquire Z-stacks. Quantification of number of particles per volume of root epidermal cells using the disector approach was applied (DIS), and maximum intensity projections were generated from the disector volumes. Autophagosomes were then counted at these maximum intensity projections, and obtained values were divided by the volumes of respective disectors (MIP). Means ± SD are presented. Pair-wise t-test, ** p < 0.01, n.s. – not significant, n = 13 – 15. Three biological replicates providing similar results were conducted independently. **C, D:** Maximum intensity projection contains autophagosomes present at the exclusion (look-up) plane of disector, root epidermal cells in the differentiation zone, autophagy-inducing treatment. **C**: Maximum intensity projection constructed from focal planes # 1 – 25 (i. e. volume of the corresponding disector without its look-up plane). **D**: Focal plane # 26 (look-up plane) of the corresponding disector. Autophagosomes excluded with the disector method but included in maximum intensity projection are marked with arrowheads. **F, G**: Particles of lower signal intensity may be overlooked in maximum intensity projections, root epidermal cells in the meristematic zone, control conditions. **E:** Maximum intensity projection constructed from focal planes # 1-25 (i. e. volume of the corresponding disector without its look-up plane). **F**: Focal plane # 12 with a particle (square) that may not be recognized at the maximum intensity projection due to its low signal. **G**: Focal plane # 18 with a particle (square) that is not apparent at the maximum intensity projection. **H, I**: Particles in the insets at **F, G** are shown at five consecutive focal planes within the disector (both are included as they disappear before the look-up plane is reached). Scale bars 10 µm.

We suppose that the observed higher values for MIP are reached in DZ because the extra particles that are included using this method and that are discarded when the disector is used are the ones that appear close to the look-up plane of the disector and which are then apparent at MIP (Fig. 5 C, D). In MZ, especially in the control conditions, the signal from the cytoplasm surrounding autophagosomes is relatively high (cf. Fig. 2 D, E). Some autophagosomes show only moderate signal intensity (Fig. 5 F, G). The same particle can, however, merge with the surrounding cytoplasm or be completely absent at the maximum intensity projections (Fig. 5 E). These are clearly apparent and recognizable, especially when observed at several focal planes, which is what is carried out using disector quantification (Fig. 5 H, I). Quantification with disector thus outperforms counting at MIP as it allows more sensitive detection of individual autophagosomes withing the sampling volume, and, on the other hand, it excludes the autophagosomes crossing disector boundaries which are erroneously included at MIP.

## DISCUSSION

We have presented the use of immunolabelling coupled with stereological quantification of autophagosomes in *A. thaliana* root. Probing with anti-ATG8 primary antibody provided a clear signal specific for developing or mature autophagosomes. Although there are a few reports on immunolabelling of cells in DZ, we demonstrated its successful application for autophagosome detection and quantification in both DZ and MZ. The fixation preceding the immunolabelling step allowed easy detection and counting of autophagosomes in 3D, which may be complicated in live imaging due to their high mobility, especially in MZ. Fixation also allows the precise termination of the treatment at a chosen timepoint, and it rules out the possibility of enhancement of autophagy arising from exposing seedlings to additional stress during sample preparation and image acquisition. Hence, it can also be an option to introduce a fixation step when the FP-ATG8 marker line is used, if convenient.

Individual isoforms of ATG8 tagged with fluorescent proteins are the most commonly used autophagosome markers. Recently, GFP-*At*ATG8a has been shown to label not only autophagosomes but also phagophores that can be detected as smaller bodies in proximity of the endoplasmic reticulum with confocal microscopy [8]. As all nine isoforms of *At*ATG8 interact with *At*SH3P2 [38], a protein required for phagophore formation and expansion [2], it is very likely that all *At*ATG8 isoforms are also localized at the phagophores. Many of the smaller particles we detected using immunolabelling may have thus been phagophores and not sealed autophagosomes. However, we did not distinguish between one type of immunolabelled particles and the other and counted both together as the sorting of FP-ATG8-decorated particles into phagophores and autophagosomes is rather rare in live imaging as well [6].

We have demonstrated that immunolabelling detects more autophagosomes than when the FP-*At*ATG8e marker is used. In addition, we showed that sub-populations of autophagosomes can be decorated with different individual isoforms of *At*ATG8. Uneven numbers of autophagosomes were observed when different *At*ATG8 isoforms fused with GFP were employed as autophagosome markers under the same treatment [8]. It was also reported that transcription response varies among single *A*tATG8 isoforms under stress [15,39,40]. All this being considered together implies that individual *At*ATG8 isoforms can be specifically employed in different biological processes. Moreover, *At*ATG8 isoforms, when used as autophagosome markers, are rarely expressed from their native promoters [15], constitutive promoters providing higher than native *AtATG8* expression levels are usually used instead [9,13,14]. Overexpression of *AtATG8* has been reported to enhance the expression of other *ATG*s, including other *ATG8* isoforms. This effect promotes the overall autophagy machinery and increases the total number of autophagosomes [16]. Using FP-*AtATG8* isoform under constitutive promoters as autophagosome markers can thus interfere with endogenous mechanisms driving autophagy. Such positive feedback can, in turn, lead to an increased number of autophagosomes when compared with an experimental set-up unaffected by such bias.

Immunolabelling with anti-ATG8 antibody does not suffer from the mentioned inconveniences as it detects a broad range of autophagosomes decorated virtually with all *At*ATG8 isoforms (possibly with the exception of *At*ATG8i) and thus provides a biologically relevant and unbiassed estimate of actual autophagosome number. Although the procedure is more demanding to perform with appropriate accuracy, it is readily available and avoids laborious and time-consuming generation of marker lines involving crossing or transformation. Therefore, immunofluorescence microscopy is a convenient and more precise alternative to the gold standard of live imaging of autophagosome using FP-ATG8 markers.

Besides FP-ATG8 markers, staining with acidotropic dyes such as monodansylcadaverine or LysoTracker was reported as a method suitable for autophagosome labelling [3,5,14]. Importantly, it has been demonstrated that particles stained with these dyes do not co-localize with autophagosomes decorated with *At*ATG8e [6] and probably represent different compartments. Thus, they seem inappropriate for autophagosome labelling in plant cells, at least at a shorter duration (up to 4 hours) of autophagy induction [41]. However, various shorter treatments taking from tens of minutes to a few hours are widely applied (recently, e.g.: [4,8–10,12,42]), and experimenters should be careful and conscious of this putative limitation of using acidotropic dyes in such cases. Autophagosomes can also be observed in differential interference contrast (DIC) setup when the number of vesicular bodies increases upon autophagy induction. For DIC observation, autophagy induction treatment is usually coupled with treatment with inhibitors preventing vacuolar degradation or fusion of autophagosomes with the vacuole [6,13,16] in order to increase the number of autophagic bodies and thus facilitate their detection. Such an approach directly applicable without introducing FP-ATG8 markers. However, small and dense particles may be hard to quantify in DIC images precisely.

Stereological methods are – unfortunately – used much less frequently in plant science than in animal or human anatomy (for a review summarizing the application of stereology in plants, see [21]), although they were reported to outperform object counting in 2D sections [30]. Optical disector combined with the Cavalieri principle has recently been applied to determine the number of chloroplasts per volume of spruce needle or in single mesophyll cells [30,43], i. e. to count individual particles within a tissue which is very similar to the quantification of autophagosomes. That is why we consider stereological quantification a perfectly suitable tool for the assessment of the number of autophagosomes.

Image analysis procedures employed to quantify autophagosomes reported in the literature are not unanimous. What is worse is that they are often only vaguely described [42]. Counting autophagosomes at individual optical 2D sections of a cell [2,3,8,44] can hardly be considered systematic. Sometimes authors use Z-stacks acquisition [6,45] which – at least in some cases – enables systematic and regular sampling of investigated cells or tissue. However, autophagosomes are then counted in separated focal planes of the Z-stack, and their counts are related to surface area units. Spacing between Z-stack focal planes is set to 1 µm in [45]. Given that such a dimension is considered the minimum diameter of autophagosomes [6], it is likely that the same autophagosome can be detected in neighbouring focal planes. Thus, their number can be overestimated if they are counted in each focal plane of a given Z-stack separately. Biased results can be obtained if this possibility is not taken into consideration and ruled out.

Some authors mention the quantification of autophagosomes at maximum intensity projections (MIP) constructed from Z-stack [17,35–37]. We were interested in whether such an approach could be used as counting at the 2D image is faster than tracking each autophagosome individually in the Z-direction when the disector is applied. Moreover, the particles can be easily counted or sorted by size, circularity etc., in an automated way, e.g., by the Analyze Particle tool in Fiji. Counting autophagosomes at MIPs constructed from the regions of interest corresponding with the volume of the disector was coupled with an estimate of the volume of the corresponding disector by the Cavalieri principle so that the resulting numbers could be related to volume and not to surface area. Interestingly we observed different results between the two methods but only in some cases, and the differences were of opposite directions based on the root zone and treatment condition.

When these observations are considered together, it can be hypothesised that different results are obtained when moderate numbers of particles per tissue volume are counted, such as in DZ under autophagy induction or in MZ in control conditions. If the total number of autophagosomes is low, the number of particles differentially detected by the two methods is also low and therefore, they do not provide sufficiently different values. On the other hand, with the very high total number of autophagosomes in the MZ during induced autophagy, the number of autophagosomes that are differentially detected by the two methods is not significant enough to cause a meaningful difference because their proportion of the total number of autophagosomes is relatively low.

In our work, we focused on the autophagy process in the root epidermis as it is directly impacted by the treatment we applied. That is why, we expected to observe the most striking effects there. To that purpose, we set up sampling using disector. Its Z-dimensions were set to 7.5 µm in to ensure it would not exceed the epidermal layer. In DZ, we chose bigger X and Y dimensions of the disector square base as we aimed to cover larger region of a root containing several cells, which were quite variable in the number of autophagosomes. In MZ, smaller X and Y dimensions were used as the cells were smaller, and the numbers of autophagosomes were relatively high, making the quantification in a greater volume too time-consuming. Two disectors were used in MZ to cover variability within this zone. Alternatively, stereological quantification of autophagosomes in individual cells coupled with the cell volume assessment using, e.g. Fakir method [46] would bring more precise information than our quantification in bulk tissue. A detailed analysis could be conducted, e.g., how different cell types, such as tricho-vs atrichoblasts in the root case, respond to autophagy-inducing treatment, however, such types of analysis was beyond the scope of our research for the time being.

We hope that our article can contribute to better appreciation and application of the stereological approach in the community of plant biologists The main drawback of applying the presented methods is their significant time requirement. While quantification in 2D can be automated as the objects can often be easily segmented from their background, such improvements still remain to be achieved in 3D. Flawless appraisal and interconnection of what is one object visualized consecutively at neighbouring Z-stack planes is probably highly demanding task to be automated and as such it is still necessary to analyse this type of data by the naked eye of a trained experimenter. Recently, deep learning employing disector has been used to quantify immunolabelled neurons in brain tissue, which performed with comparable accuracy but higher efficiency than human [47,48]. It can be thus assumed that introduction of artificial intelligence in image analysis will enormously enhance the use and performance of stereological methods; a decrease in the time required for analysis spent by a human experimenter is expected. Hopefully, this will enable stereological approaches to be more employed in the field of plant anatomy and cell biology, contributing to higher accuracy and better reproducibility of quantitative data analyses. Autophagy studies can significantly benefit from such development as autophagosome detection and counting are essential ways to study such a vital process.

## ABBREVIATIONS

AF-555: Alexa Fluor 555

ATG: autophagy related

ATG8: autophagy related 8

*At*ATG8a-i: autophagy related 8 from *Arabidopsis thaliana*, isoforms

*At*TG8a to *At*ATG8i *atg10-1*: *A. thaliana* loss of function mutant line of autophagy related 10

BSA: bovine serum albumin

DIS: disector

DL-488: DyLight 488

DMSO: dimethyl sulfoxide

DTT: dithiothreitol

DZ: differentiation zone (of the root)

EGTA: 3,12-Bis(carboxymethyl)-6,9-dioxa-3,12-diazatetradecane-1,14-dioic acid

estV: estimate of tissue volume

estNV: estimate of number of particles per tissue volume

FP-ATG8: autophagy related 8 protein fused to a fluorescent protein

GFP: green fluorescent protein

MES: 2-(morpholin-4-yl)ethanesulfonic acid

MIP: maximum intensity projection

MTSB: microtubule stabilizing buffer

MTSB-T: microtubule stabilizing buffer containing Triton X-100

MZ: meristematic zone (of the root)

PAR: photosynthetically active radiation

PFA: paraformaldehyde

PIPES: 2,2′-(Piperazine-1,4-diyl)di(ethane-1-sulfonic acid)

## ACKNOWLEDGEMENTS

The project was supported by Horizon 2020 COST action TRANSAUTOPHAGY (CA15138) – grant No. LTC17084, and by European Regional Development Fund—Project “Centre for Experimental Plant Biology” No. CZ.02.1.01/0.0/0.0/16_019/0000738; both Ministry of Education, Youth and Sports of the Czech Republic. The Imaging Facility of the Institute of Experimental Botany was supported by “National Infrastructure for Biological and Medical Imaging (Czech-BioImaging – LM2018129)” project, and by European Regional Development Fund (Project No. CZ.02.1.01/0.0/0.0/18_046/0016045); both Ministry of Education, Youth and Sports of the Czech Republic.

The authors would like to thank Dr Barbora Radochová, Institute of Physiology, Prague, Czech Republic, for consulting and helping with the stereological methods; to Dr Jan Petrášek, Dr Adriana Jelínková and Karolína Holečková, all Institute of Experimental Botany, Prague, Czech Republic; for the introduction to immunolabelling procedure and occasional assistance with the automated pipetting platform, to Dr Yasin Dagdas, Gregor Mendel Institute of Molecular Plant Biology, Vienna, Austria for sharing *A. tumafaciens* strains harbouring GFP-*At*ATG8a-i constructs.

## SUPPLEMENTAL FIGURES

**Figure S1:**
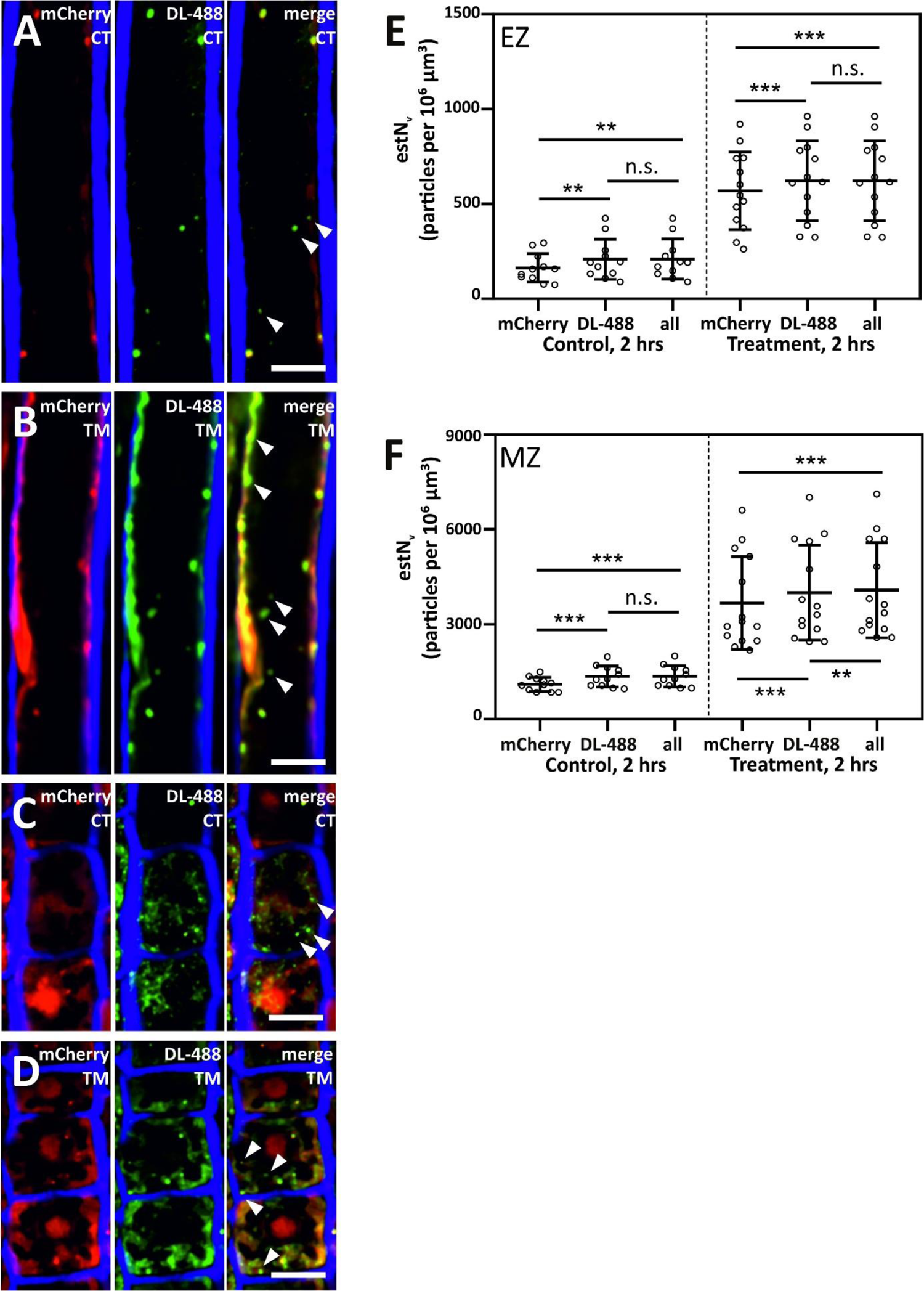
Anti-ATG8-labelled autophagosomes colocalize with mCherry-ATG8e. **A – D:** Colocalization of autophagosomes labelled with mCherry-*At*ATG8e (left panel, in red) with the immunolabelled ones (middle panel, in green) in root epidermal cells in the differentiation (**A, B**) or meristematic zone (**C, D**) under control conditions (CT; **A, C**) or under the treatment inducing autophagy (TM; **B, D)**. Immunolabelled autophagosomes devoid of mCherry signal are observed (arrowhead in merge images, right panel) whereas the majority of particles show clear signal in both channels. Roots of 6-day-old seedlings expressing 35S::mCherry-ATG8e were immunolabelled, primary antibody: anti-ATG8, secondary antibody: anti-rabbit DyLight 488, and subsequently counter-stained with CalcoFluor White to visualize cell borders (in blue). Spinning-disk microscope was used to acquire Z-stacks. Maximum intensity projections of six Z-planes (i. e. 1.5 µm depth along the Z-axis) are presented. Scale bars: 10 µm. **E, F:** Stereological quantification of number of particles per volume of root epidermal cells in differentiation zone (DZ) and meristematic zone (MZ). Particles were counted separately in individual channels (mCherry: sum of particles detected with mCherry-*At*ATG8e; DL-488: sum of particles detected with anti-ATG8/anti-rabbit DyLight488; all: sum of all detected particles). Means ± SD are presented. Pair-wise t-test, ** p < 0.01; *** p < 0.001, n. s. not significant. n = 10 – 16. Two biological replicates providing the same results were conducted independently.

**Figure S2:**
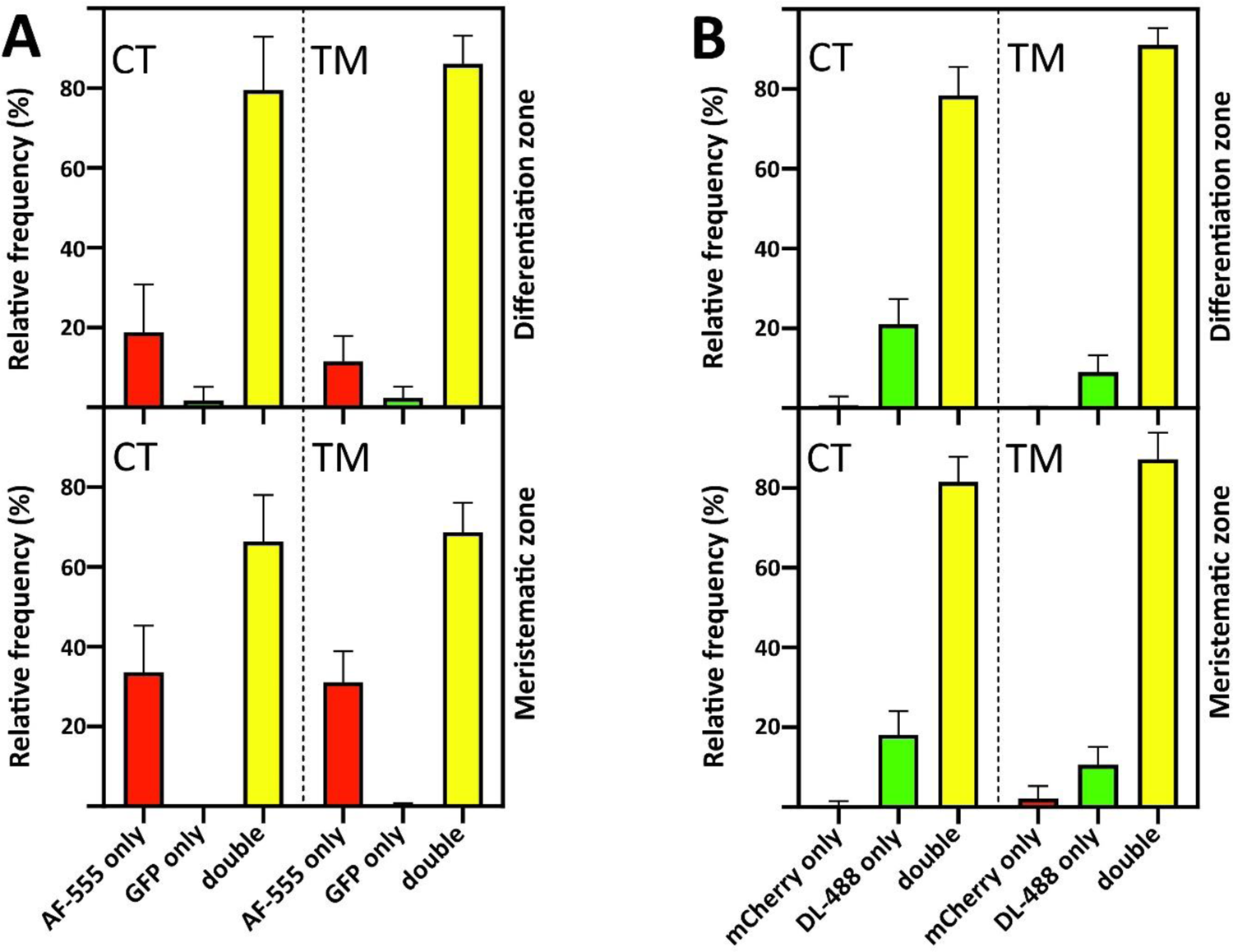
Distribution of co-localizing and non-co-localizing autophagosomes in roots of seedlings expressing 35S::GFP-ATG8e immunolabelled with anti-ATG8 and anti-rabbit AlexaFluor 555. (**A**) or 35S::mCherry-ATG8e immunolabelled with anti-ATG8 and anti-rabbit DyLight 488 (**B**) under control conditions (CT) and under treatment inducing autophagy (TM) presented in Figure 2 and Figure S3. AF-555 only, and DL-488 only: autophagosomes showing exclusively immunolabelling signal. GFP only, and mCherry only: autophagosomes showing exclusively signal of *At*ATG8e fused to the respective fluorescent protein. Double: autophagosomes with the co-localizing signal of immunolabelling and *At*ATG8e fused to the respective fluorescent protein. Histograms of means ± SD are presented. Three (A) and two (B) biological replicates providing the same results were conducted independently.

**Figure S3:**
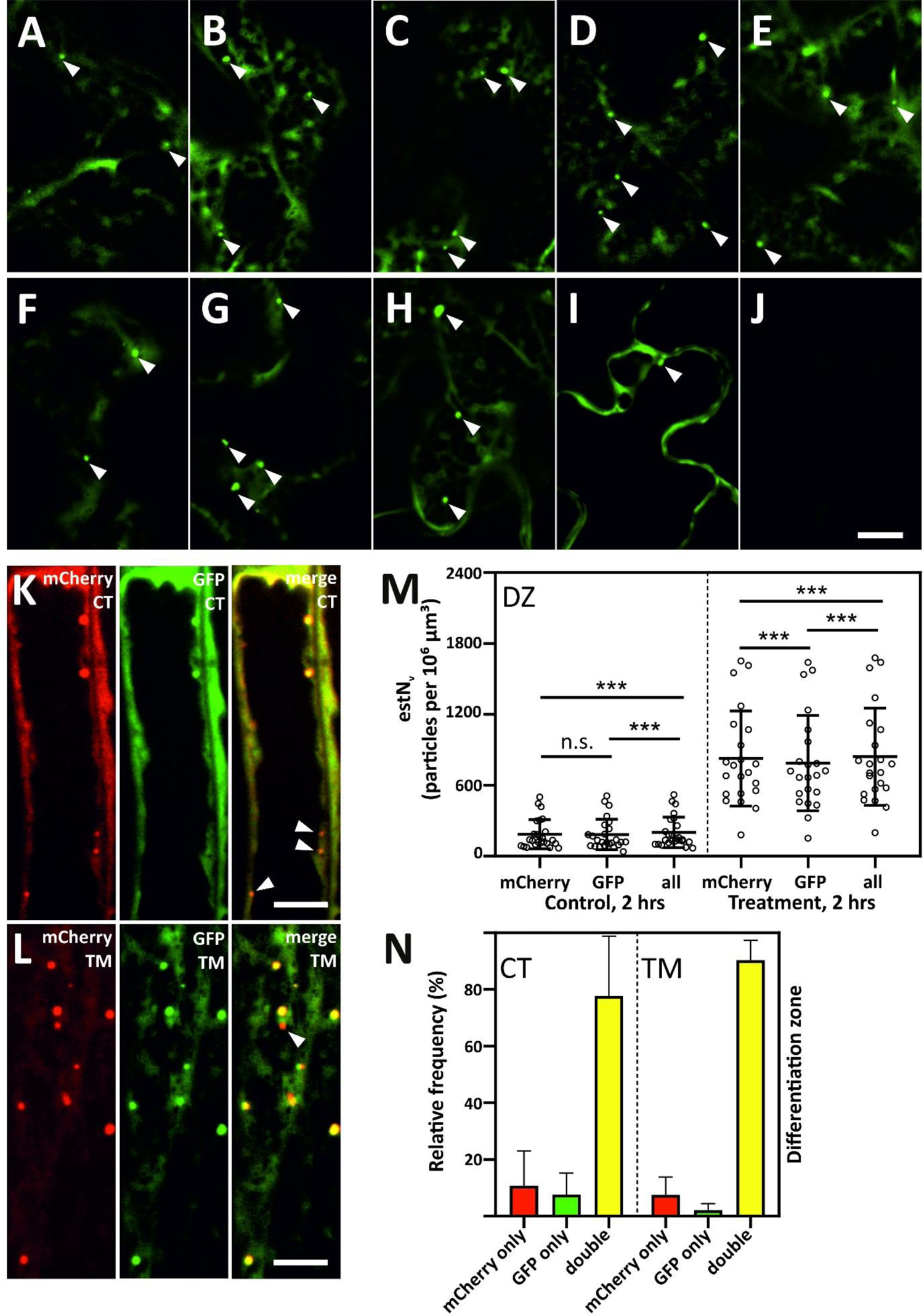
**A – J**: Transient expression of *At*ATG8 isoforms tagged with GFP in *N. benthamiana* leaf epidermal cells. **A:** GFP-*At*ATG8a. **B:** GFP-*At*ATG8b. **C:** GFP-*At*ATG8c. **D:** GFP-*At*ATG8d. **E:** GFP-*At*ATG8e. **F:** GFP-*At*ATG8f. **G:** GFP-*At*ATG8g. **H:** GFP-*At*ATG8h. **I:** GFP-*At*ATG8i. **J:** p19 only. Putative autophagosomes are marked with arrowheads. **K – M:** Different isoforms of *At*ATG8 label specific subpopulations of autophagosomes in root epidermal cells in the differentiation zone of a marker line co-expressing 35S::GFP-*At*ATG8a and 35S::mCherry-*At*ATG8e. **K**: control conditions (CT). **L**: treatment inducing autophagy (TM). **K**, **L**: left panel: mCherry-*At*ATG8e, middle panel: GFP-*At*ATG8a, right panel: merge; arrowheads mark non-co-localizing autophagosomes. Images represent maximum intensity projections of six Z-planes (i. e. 1.5 µm depth along the Z-axis). Scale bars: 10 µm. **M**: Stereological quantification of number of particles per volume of root epidermal cells in differentiation zone (DZ). Particles were counted separately in individual channels (mCherry: sum of particles detected with mCherry-*At*ATG8e; GFP: sum of particles detected with GFP-*At*ATG8a; all: sum of all detected particles). Means ± SD are presented. Pair-wise t-test, *** p < 0.001, n. s. not significant, n = 22 – 25. **N**: Distribution of co-localizing and non-co-localizing autophagosomes in roots of seedlings co-expressing GFP-*At*ATG8a and 35S::mCherry-ATG8e under control conditions (CT) and under treatment inducing autophagy (TM). mCherry only: autophagosomes showing exclusively mCherry signal. GFP only: autophagosomes showing exclusively GFP signal. Double: autophagosomes with co-localization of both signals. Histograms of means ± SD are presented. Two biological replicates providing similar results were conducted independently.

**Figure S4:**
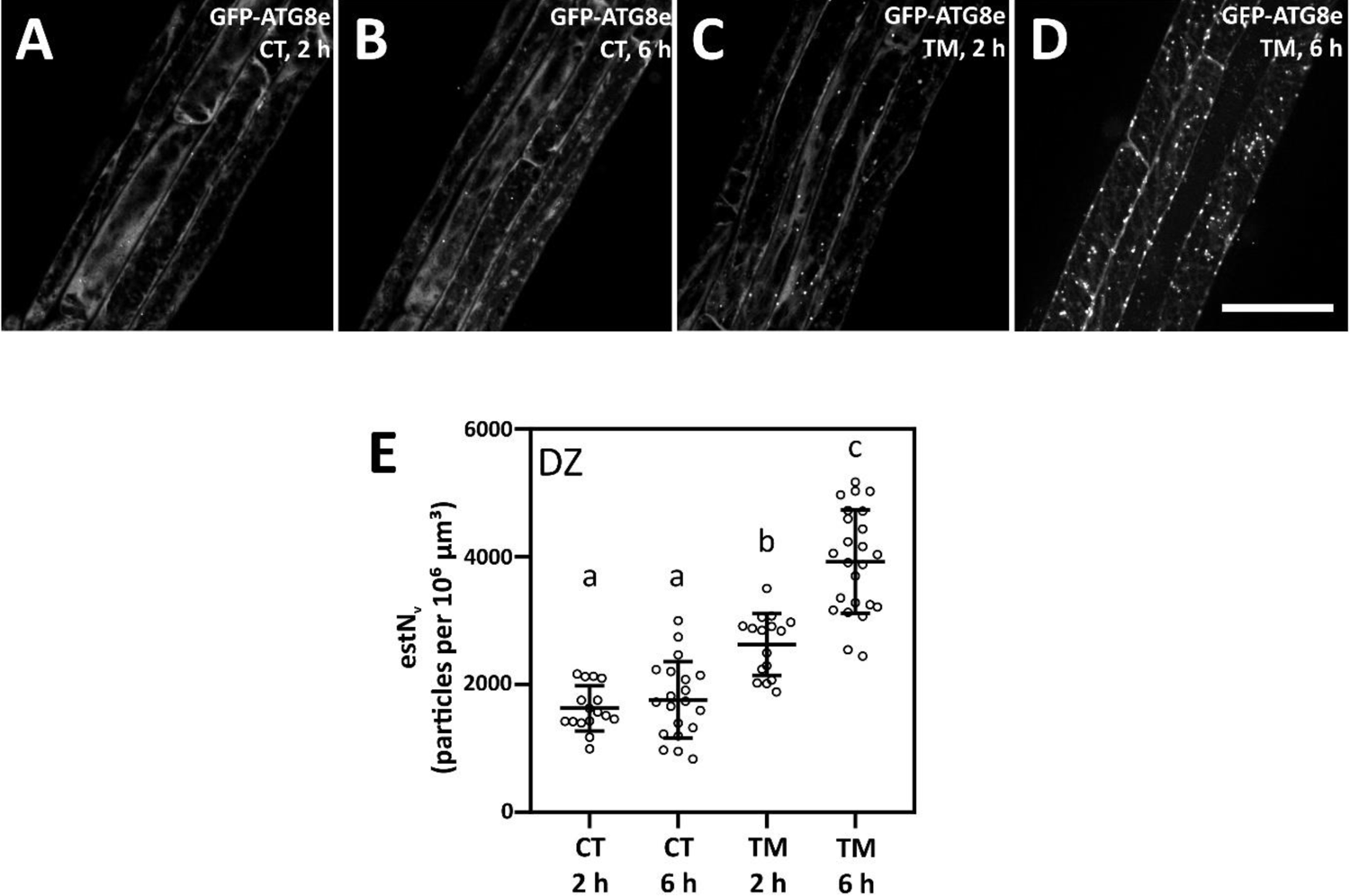
Longer duration of autophagy-inducing treatment increases the formation of autophagosomes in root epidermal cells in the differentiation zone of the marker line expressing 35S::GFP-*At*ATG8e. **A** – **D**: Seedlings grown in control conditions were transplanted to the same conditions (CT) for 2 hours (**A**) and 6 hours (**B**) or to the conditions inducing autophagy (TM) for 2 hours (**C**) and 6 hours (**D**). Spinning-disk microscope was used to acquire Z-stacks. Images represent maximum intensity projections of disector volumes used for stereological quantification. Scale bar: 50 µm. **E**: Quantification of number of autophagosomes per volume of tissue in root epidermal cells in differentiation zone under control and autophagy inducing treatment applied for 2 hours or 6 hours. Means ± SD are presented. One-way ANOVA, letters indicate significant differences between groups by multiple comparison Tukey-Kramer test, p < 0.01, n = 16 – 24.

