## Supplementary figures and images for "A novel workflow for unbiased quantification of autophagosomes in 3D in *Arabidopsis thaliana* roots"

### Fig. S1

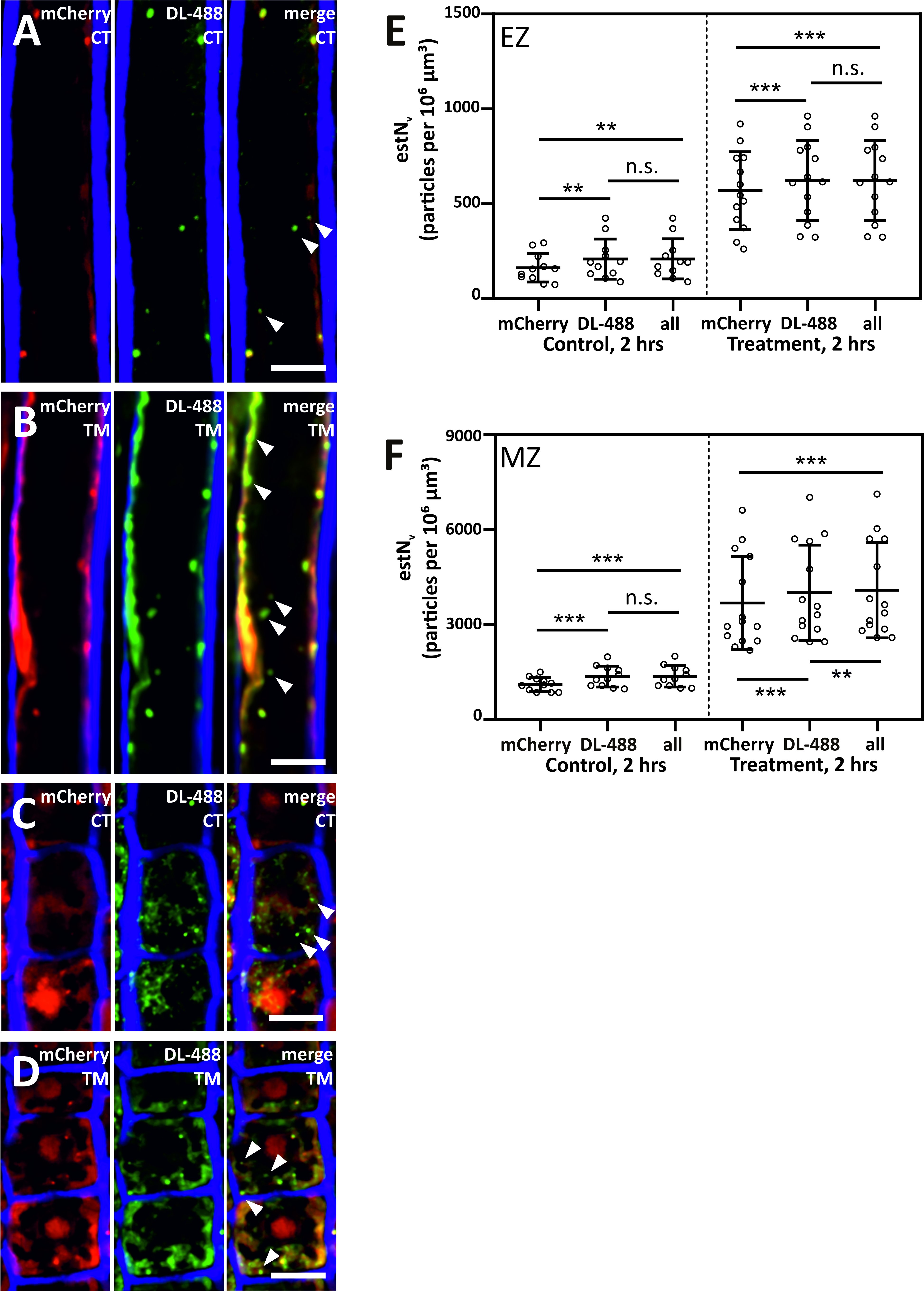

### Fig. S2

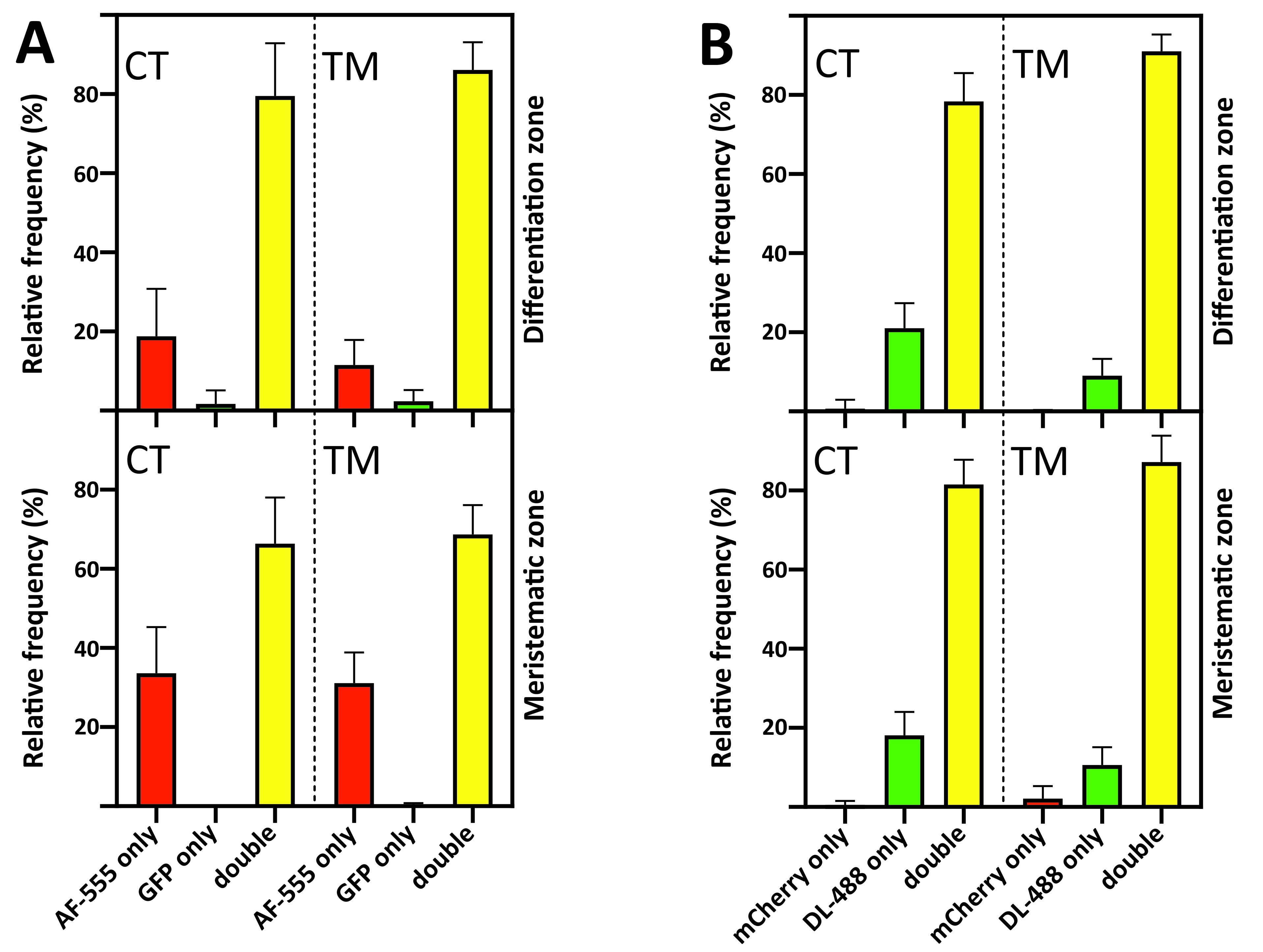

### Fig. S3

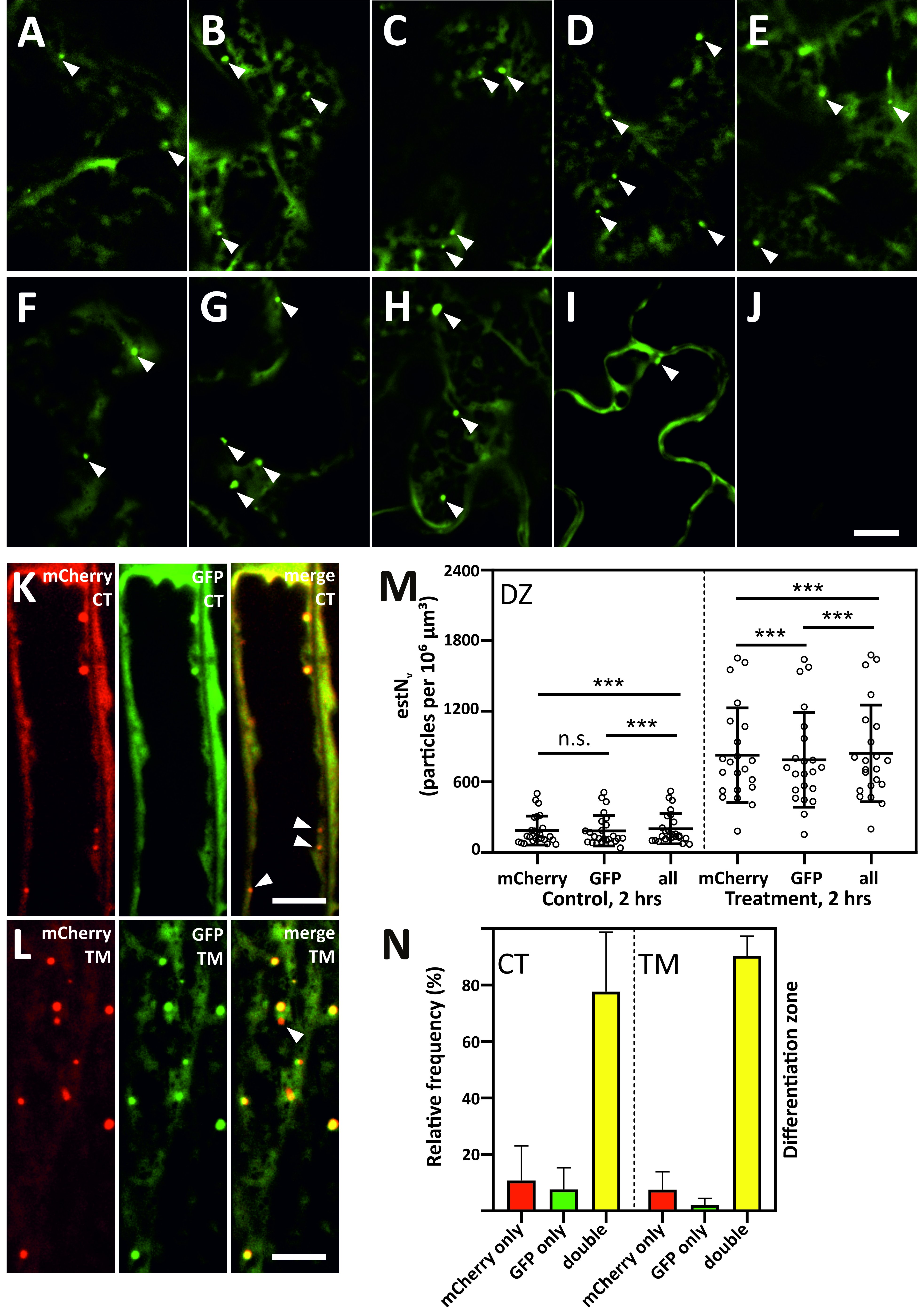

### Fig. S4

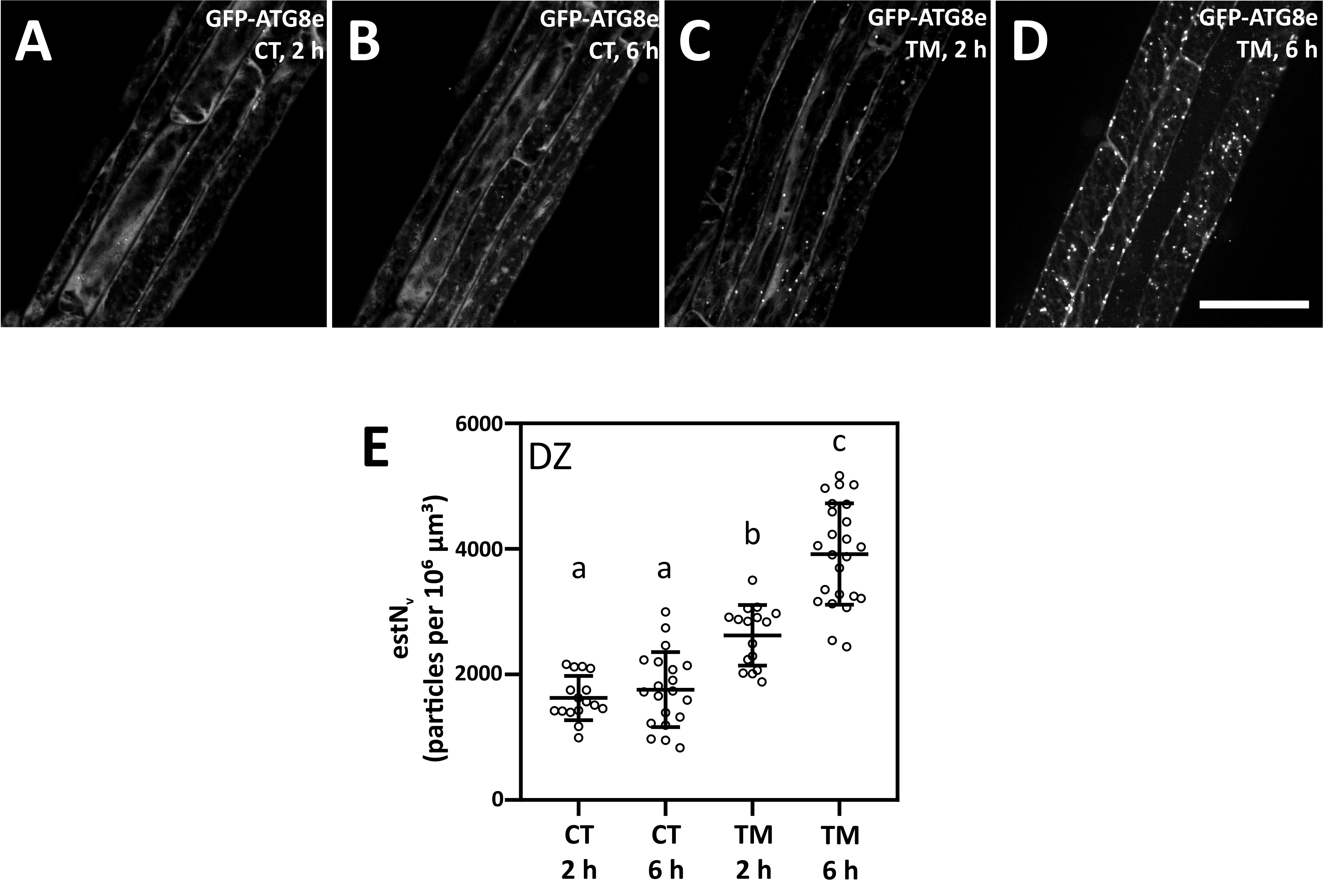
