## Supplement 1 for "A novel workflow for unbiased quantification of autophagosomes in 3D in *Arabidopsis thaliana* roots"

### Anti-ATG8 immunolabelling protocol for *Arabidopsis thaliana* roots

**Material and Equipment**

- Microscope slides and coverslips (e.g. Marienfeld 1000000 and 0101192). Slides are washed thoroughly with distilled water with detergent and in pure ethanol prior to use.
- Parafilm
- Incubation baskets (Intavis 12.440)
- 0.22 µm syringe filter (Merck SLGS033SS)
- Automated pipetting station (Intavis InSituPro VS)
- Dissecting microscope
- Dissecting scissors
- Fine tweezers
- 24-well cell culture plates

**Reagents**

- anti-ATG8 primary antibody (Agrisera; AS142769)
- anti-rabbit-DyLight488 secondary antibody (Agrisera; AS09633)
- BSA (Albumin fraction V (from bovine serum); Merck 112018)
- CalcoFluor White (Fluorescent Brightener 28 disodium salt; Sigma 910090)
- DMSO (dimethyl sulfoxide; WWR 85396.290E)
- EGTA (3,12-Bis(carboxymethyl)-6,9-dioxa-3,12-diazatetradecane-1,14-dioic acid; Sigma 03777)
- Glycerol (MP Biomedicals 800688)
- Igepal CA-630 (octylphenoxy poly(ethyleneoxy)ethanol; Sigma I8896)
- MgSO_4_.7H_2_O (magnesium sulphate heptahydrate; Merck 105886)
- Pectolyase (pectolyase Y-23 from *Aspergillus japonicus*; Duchefa Biochemie P8004)
- PFA (paraformaldehyde; Sigma P6148)
- PIPES (2,2′-(piperazine-1,4-diyl)di(ethane-1-sulfonic acid); Sigma P6753)
- Triton X-100 (2-[4-(2,4,4-trimethylpentan-2-yl)phenoxy]ethanol; MP Biomedicals 807426)
- ddH_2_O or Milli-Q H_2_O

**Steps in the Procedure**

Step 1: **Microtubule-stabilizing buffer preparation**

- Microtubule-stabilizing buffer (MTSB) is used throughout the procedure. Composition: 50 mM PIPES, 5 mM EGTA, 5 mM MgSO_4_.7H_2_O, pH = 6.8 (5M KOH). Add Triton X-100 to 0.1 % final concentration (v/v) to obtain MTSB-T.

Note: PIPES is an acid and will not dissolve in water unless the solution is alkalized.

Step 2: **Fixation**

- PFA is depolymerized in MTSB-T by heating the solution to 75 °C in a covered container in a fume hood to obtain a final concentration of 4 % (w/v). It is let cool down to the laboratory temperature. If the solution volume is reduced due to evaporation during heating, the original volume is topped up with water. The solution is filtered through 0.22 µm syringe filter to remove traces of undepolymerized PFA.
- 1 mL of fixation solution is pipetted in wells of the 24-well plate. The incubation baskets are placed in the wells. Treated seedlings (whole seedlings can be used up to 4 days of age) or detached root tips of approximately 2 cm (in the case of older seedlings) are put in the incubation baskets. Six whole seedlings or 8-10 detached roots can fit into one basket.
- Fixation is carried out for 1 hour at room temperature. To terminate fixation the baskets are moved to wells of the Intavis specimen holder (equipment of the InSituPro VS station), each containing 600 µL of MTSB-T.

The specimen holder is mounted in the automated station and the steps 3 – 9 are carried out in the station. The volume of each solution pipetted per one incubation basket is 600 µL.

Step 3: **Initial rinse and equilibration**

- 15 mins with MTSB-T, room temperature; 5 times
- 15 mins with 0.1 % Triton X-100 in water, room temperature; 5 times

Step 4: **Cell wall digestion** (+ wash)

- 30 mins with 0.05 % pectolyase in MTSB-T (w/v), 37 °C
- 15 mins with MTSB-T, room temperature; 5 times

Step 5: **Membrane permeabilization** (+ wash)

- 30 mins with 10 % DMSO + 3 % Igepal CA-630 in MTSB-T (v/v), room temperature; twice
- 15 mins with MTSB-T, room temperature; 5 times

Step 6: **Blocking**

- 1 hour with 2 % BSA in MTSB-T (w/v), room temperature

Step 7: **Primary antibody incubation** (+ wash)

- 6 hours with anti-ATG8 diluted 1:1000 in 2 % BSA in MTSB-T (filtered through 0.22 µm syringe filter to remove aggregates), 37 °C
- 15 mins with MTSB-T, room temperature; 8 times

Note: primary antibody can be re-used a few times

Step 8: **Secondary antibody incubation** (+ wash)

- 5 hours with anti-rabbit-DyLight488 diluted 1:1000 in 2 % BSA in MTSB-T (filtered through 0.22 µm syringe filter to remove aggregates), 37 °C
- 15 mins with MTSB-T, room temperature; 5 times
- 15 mins with water, room temperature; 5 times

Step 9: **CalcoFluor White counterstaining** (+ final wash)

- 20 mins with 0.002 % CalcoFluor White in water, room temperature
- 15 mins with water, room temperature; 8 times

Specimens in baskets are removed from the pipetting station and put in wells of a 24-well plate each containing 1 mL 50 % glycerol (v/v)

Step 10: **Specimen mounting**

- Strips of one layer of Parafilm are pressed onto microscopy slides in order to prevent direct contact between the slide and coverslip and to avoid mechanical damage to the roots. Immunolabelled roots are carefully removed from the incubation baskets using tweezers and mounted on slides. A dissecting microscope may be necessary to locate the roots in the baskets. 50 % glycerol (v/v) is used as the mounting medium. Sodium azide (0.02 %) may be added to the mounting medium for long-term storage of specimens (several months) in the cold.
